# An Autophagy-Dependent Tubular Lysosomal Network Synchronizes Degradative Activity Required for Muscle Remodeling

**DOI:** 10.1101/2020.04.15.043075

**Authors:** Tadayoshi Murakawa, Amy A. Kiger, Yuriko Sakamaki, Mitsunori Fukuda, Naonobu Fujita

**Affiliations:** Cell Biology Center, Institute of Innovative Research, Tokyo Institute of Technology, 4259-S2-11 Nagatsuta-cho, Midori-ku, Yokohama, Kanagawa 226-8503, Japan; Laboratory of Membrane Trafficking Mechanisms, Department of Integrative Life Sciences, Graduate School of Life Sciences, Tohoku University, Aoba-ku, Sendai, Miyagi 980-8578, Japan; Section of Cell and Developmental Biology, Division of Biological Sciences, University of California, San Diego, La Jolla, CA, 92093, USA; Microscopy Research Support Unit Research Core, Tokyo Medical and Dental University, Tokyo, 113-8510, Japan; Precursory Research for Embryonic Science & Technology (PRESTO), Japan Science & Technology Agency (JST), 4-1-8 Honcho Kawaguchi, Saitama, 332-0012, Japan

**Keywords:** autophagy, muscle, tubular lysosome, autolysosome, Drosophila, metamorphosis, myofiber, atrophy, Syntaxin17

## Abstract

Previously, we reported that autophagy is critical for Drosophila muscle remodeling during metamorphosis (Fujita et al., 2017). However, little is known about how lysosomes meet increased degradative demand upon cellular remodeling. Here, we found an extensive tubular autolysosomal network in remodeling muscle. The tubular network transiently appeared and exhibited the capacity to degrade autophagic cargoes. The tubular autolysosomal network was uniquely marked by the autophagic SNARE protein, Syntaxin 17, and its formation depended on both autophagic flux and degradative function, with the exception of the Atg12 and Atg8 ubiquitin-like conjugation systems. Among *ATG*-deficient mutants, the efficiency of lysosomal tubulation correlated with the phenotypic severity in muscle remodeling. The lumen of the tubular network was continuous and homogeneous across a broad region of the remodeling muscle. Altogether, we revealed that the dynamic expansion of a tubular autolysosomal network synchronizes the abundant degradative activity required for developmentally regulated muscle remodeling.

**Impact Statement:** Analysis of developmentally-regulated Drosophila muscle remodeling revealed autophagy-dependent formation of an extensive, Syntaxin 17-marked, tubular network that synchronizes the abundant degradative activity across a broad region of the remodeling muscle

## Introduction

Lysosomes are membrane-bound compartments for the degradation of both endocytic and autophagic cargoes in the eukaryotic cell. The lumen of lysosomes maintains an acidic pH to digest materials by a series of acid hydrolases (Lawrence and Zoncu, 2019). In addition to the catabolic function, lysosomes play numerous roles, such as secretion, nutrient sensing, and signaling through mechanistic target of rapamycin (mTOR) complex I and AMP-activated protein kinase (AMPK). Thus, the regulation of lysosomal function is critical for cellular homeostasis. The MiT/TFE family of transcription factors, including TFEB and TFE3, are master regulators of the expression of a myriad of lysosomal and autophagic functions needed to meet changing degradative demands (Martina et al., 2014; Sardiello et al., 2009). However, very little is known about the mechanisms that mediate the modulation of lysosomal degradative capacity through coordinated changes in activity, quantity, distribution, and morphology (Hipolito et al., 2018).

Although lysosomes are generally thought of as spherical organelles, lysosomal shape undergoes morphological changes in response to certain conditions. The existence of tubulated lysosomes, called tubular lysosomes or nematolysosomes, have been known since the 1970s in various cell types, including macrophages, pancreatic exocrine cells, neurons, and muscle cells (Knapp and Swanson, 1990; Swanson et al., 1987; Okada et al., 1986; Robinson et al., 1986; Shi et al., 1992; Araki et al., 1993; Oliver, 1980). Tubulated lysosomes are prominent in lipopolysaccharide (LPS)-activated macrophages and dendritic cells (Hipolito et al., 2018). Recently, the tubular lysosomal network was described in the fly larval body wall muscle (Johnson et al., 2015) and the nematode epidermis during molting (Miao et al., 2020). In general, the extended tubular lysosomes exhibited features of typical functional lysosomes, including the accumulation of acid phosphatases, lysosomal proteases, Lamp, and vacuolar H^+^-ATPase (V-ATPase). Microtubules may template the tubulation. In the model, plus-end-directed kinesin motors and minus-end-directed dynein-dynactin complexes extend lysosomal tubules in opposite directions via functions of the Arl8b-SKIP complex and Rab7-RILP or -FYCO1 complex, respectively (Mrakovic et al., 2012). However, the stretching model alone cannot explain how the tubular lysosome becomes over 10 μm in length. To date, the mechanisms shaping tubular lysosomes are poorly understood, and the physiological significance of the tubulation remains enigmatic.

Autophagy is an intracellular bulk degradation system in which double membrane-bound autophagosomes sequester and deliver cytosolic materials to the lysosomes/vacuoles for degradation. Autophagosome formation is mediated by at least 18 core autophagy-related (Atg) proteins acting within six functional units (Mizushima et al., 2011): 1) the ULK/Atg1 protein kinase complex; 2) the autophagy-specific phosphatidylinositol 3-kinase (PI3K) complex; 3) the phosphatidylinositol 3-phosphate (PI3P)-binding protein complex; 4) Atg9; 5) the LC3/Atg8 conjugation system; and 6) the Atg12 conjugation system. All six units are pivotal for autophagy. However, the Atg8 and Atg12 ubiquitin-like conjugation systems seem to be dispensable for the elongation of the autophagic membrane in mammalian cells (Tsuboyama et al., 2016). The completed autophagosomes then fuse with lysosomes to form autolysosomes, the site for autophagic degradation and subsequent macromolecule efflux. The fusion is mediated by two soluble NSF attachment protein receptor (SNARE) complexes, Syntaxin17 (Qa)-SNAP29 (Qbc)-VAMP8 (R) and Syntaxin7 (Qa)-SNAP29 (Qbc)-Ykt6 (R), (Itakura et al., 2012; Takáts et al., 2013; Matsui et al., 2018; Takáts et al., 2018). It has been reported that autolysosomes tubulate in the process of lysosome reformation from the autolysosome, called autophagic lysosome reformation (ALR) (Yu et al., 2010). The ALR tubule seems to be a kind of tubular lysosome; however, the tubule is neither acidic nor contains acidic hydrolases. Accordingly, it is thought of as a proto-lysosome (Yu et al., 2010).

Differentiated muscle cells, or myofibers, have highly organized and specialized organelles needed for muscle contraction. The contractile system is made up from sarcomeres arrayed into myofibrils. Sarcomere contractions are coordinated by changing levels of cytoplasmic calcium in response to a signal relay along the ‘excitation-contraction coupling’ system: transverse (T)-tubule invaginations of the plasma membrane in junctions with the sarcoplasmic reticulum. Mechanisms must remodel these organelles with ongoing muscle reorganization in response to muscle cell growth, use, damage, atrophy and aging. However, the mechanisms of muscle remodeling remain mostly unknown, in part due to challenges with observing the organellar dynamics within intact muscles. In Drosophila, a set of larval body wall muscles persist throughout metamorphosis as pupal abdominal muscles, called dorsal internal oblique muscles (DIOMs). In the DIOMs, the entire contractile and excitation-contraction coupling systems undergo a developmentally-programmed remodeling during metamorphosis (Fujita et al., 2017), providing an excellent experimental model to study mechanisms of synchronous muscle atrophy to hypertrophy (Kuleesha et al., 2016). We recently reported that autophagy plays a critical role in DIOM remodeling (Fujita et al., 2017). Upon disassembly of myofiber organization in DIOMs, the cytoplasmic contents including mitochondria were enwrapped by autophagic membranes and delivered into lysosomes for degradation. In the process, not only autophagosome formation but also lysosomal functions must be regulated. However, little is known about the mechanisms that coordinate lysosomal function with cellular remodeling.

Here, we found an extensive, tubular autolysosomal (tAL) network in Drosophila muscles during metamorphosis. The induction of autophagy with muscle remodeling was necessary for autolysosomal tubulation, which was uniquely marked by the autophagy-related SNARE, Syntaxin 17 (Stx17). The tubular network was continuous with a homogeneous lumen, and the tAL network extended over a wider range of the remodeling muscle cell than were spherical lysosomes found in the stable myofibers prior to remodeling. We show that the tubular autolysosomal network acts to synchronize activity and meet increased degradative demand with muscle remodeling.

## Results

### Syntaxin 17 marks aa tubular network in remodeling muscle cells

To gain insight into organelle dynamics with muscle remodeling during metamorphosis (Figure 1A–B), the localizations of GFP-tagged reporters for different cellular membrane compartments were observed every day after pupal formation (APF) in intact abdominal DIOMs. At the third instar larval (3IL) stage, GFP-fused Stx17, a marker of autophagic membranes in differentiated muscle (Fujita et al., 2017), was detected as previously described as vesicular structures. Strikingly, upon metamorphosis we found that GFP:Stx17 appeared as a tubular network present in all DIOMs by 1 d APF (Figure 1C). Thereafter, the extent of tubulation gradually decreased, and a GFP:Stx17 vesicular pattern was restored by 3 d APF (Figure 1C). To observe how the network developed, we performed time-course microscopy. GFP:Stx17 remained in puncta at 12 h APF, then became increasingly tubulated until 16-20 h APF (Figure 1D–E), indicating a dynamic rearrangement of GFP:Stx17-labeled membranes with the onset of DIOM muscle remodeling (Fujita et al., 2017).

**Figure 1.**
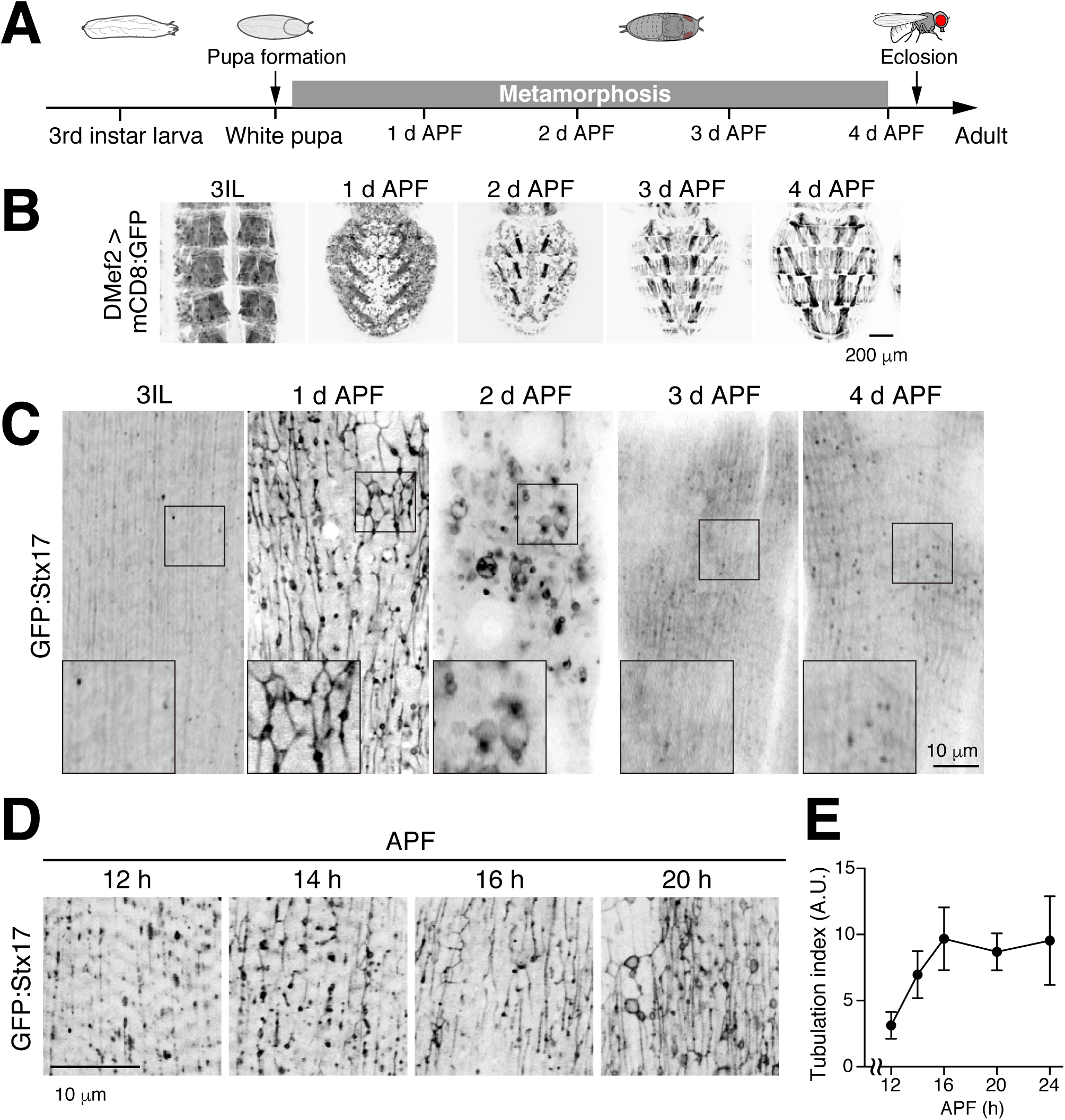
Syntaxin 17 marks a tubular network in remodeling muscle cells. (A) Timeline of fly development from third instar larva to adult at 25°C, days after puparium formation (d APF). (B) Time course microscopy of mCD8:GFP in dorsal abdominal muscles during metamorphosis. (C and D) Time course microscopy of GFP:Stx17 in dorsal internal oblique muscle (DIOM) imaged through the cuticle of live wildtype animals from third instar larvae (3IL) to 4 d APF (C), from 12 h to 20 h APF (D). (E) Quantification of GFP:Stx17-positive tubules in DIOMs from 12 to 24 h APF. The values are the mean ± standard deviation (SD), n=7.

### The Syntaxin 17 tubular network has characteristics of autolysosomes

Stx17, a SNARE protein, localizes to the autophagosome and detaches just after fusion with the lysosome (Itakura et al., 2012; Takáts et al., 2013). We have reported that autophagy is robustly induced with DIOM remodeling by 1 d APF (Fujita et al., 2017). Therefore, we postulated that the tubular network was an autophagy-related structure. To test the possibility, we performed time-course microscopy of autophagic flux over DIOM remodeling. Using the mCherry:GFP:Atg8 reporter, autophagosomes and autolysosomes can be distinguished by GFP-sensitivity to low pH (Kimura et al., 2007). While the mCherry:GFP:Atg8 signal in the GFP channel did not exhibit any tubular structures, the mCherry was distributed to highly tubulated structures (Figure 2A) that colocalized with GFP:Stx17 (Figure 2B) in 1 d APF DIOMs. This result indicates that the Stx17-positive tubular network is an autolysosome-related organelle.

**Figure 2.**
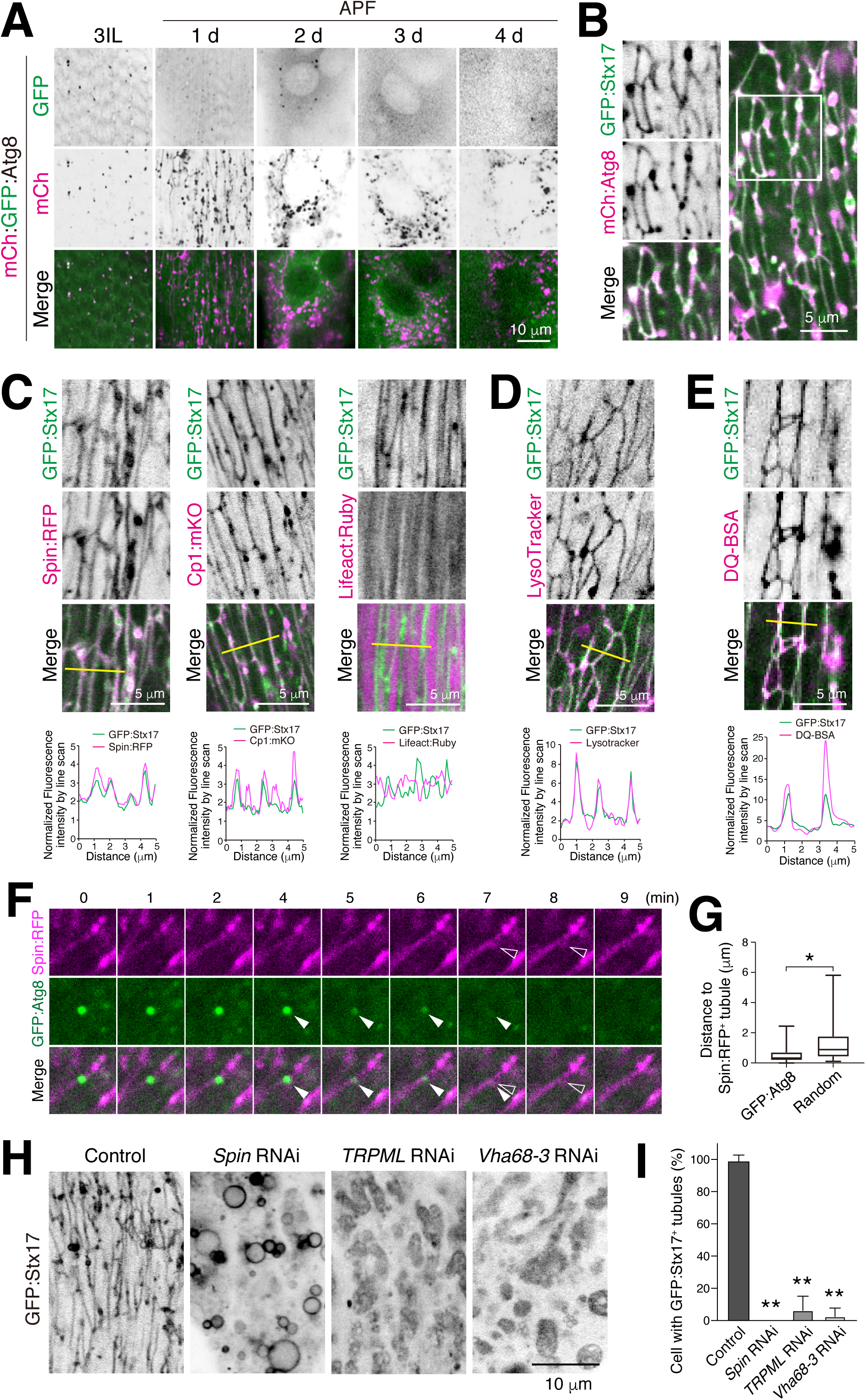
The Syntaxin 17 tubular network has characteristics of autolysosomes. (A) Time course microscopy of mCherry:GFP:Atg8 in DIOM imaged through the cuticle of live animals from 3IL to 4 d APF. (B) Colocalization between GFP:Stx17 and mCherry:Atg8 in 20 h APF DIOM. (C) Colocalization between GFP:Stx17 and Spin:RFP, Cp1:mKO, or Lifeact:Ruby in 20 h APF DIOMs. Line plot profiles of the yellow line in each panel. (D and E) Live pupae were injected with LysoTracker Red (D) or DQ Red-BSA (E), and DIOMs were imaged through the cuticle at 20 h APF. Line plot profiles of the yellow line in each panel. (F and G) Time-lapse imaging of Spin:RFP and GFP:Atg8a in 20 h APF DIOMs (F). Disappearing GFP punctum, white arrowhead; shape-change of tubular lysosome, colorless arrowhead. Quantification of the distance of Spin:RFP-positive tubule, and the point at which GFP:Atg8 puncta disappeared (GFP:Atg8) or randomly drawn puncta (random), n>30. *, p<0.05 (Student’s *t*-test) (G). (H and I) GFP:Stx17 localization in DIOMs at 20 h APF in control, *Spin* RNAi, *TRPML* RNAi, or *Vha68-3* RNAi (H). Mean percent + SD of DIOMs with more than 5 μm GFP:Stx17-positive tubules, n=10. **, p<0.001 (Dunnett’s test) (I).

We characterized the tubular compartment further. GFP:Stx17-positive tubular structures colocalized with two lysosomal proteins, Spinster, a lysosomal sugar transporter (Spin:RFP), and the cathepsin L cysteine protease (Cp1:mKO), but not with F-actin sarcomeres (Lifeact:Ruby) (Figure 2C). Moreover, compartmental acidification and degradative activity — including throughout the tubules — was indicated by GFP:Stx17 colocalization with LysoTracker Red, a dye for acidic organelles, and dye-quenched (DQ) Red-BSA, a fluorogenic substrate for proteases, respectively (Figure 2D–E). The autolysosomal activity seen within the GFP:Stx17-positive tubules contrasts the lack of activity described for tubules involved in autophagic lysosomal reformation (ALR) (Yu et al., 2010; Chen and Yu, 2013). To determine whether the autolysosomal tubules have a capacity to receive and degrade transported materials, we performed live imaging of autophagosomes (GFP:Atg8) and the autolysosome network (Spin:RFP), respectively. We observed initially bright GFP:Atg8 puncta that, over a few minutes, quenched at the site of Spin:RFP-positive tubules (Figure 2F, Figure 2—figure supplement 1, and Supplementary video 1). Concomitant with quenching of GFP:Atg8, the shape of the Spin:RFP-positive tubule was transiently distended, indicating autophagosome fusion with the tubule. We noticed that signal from GFP:Atg8 puncta disappeared near tubules in most cases. Collectively, the distances between GFP:Atg8 puncta to the nearest Spin:RFP tubule was significantly lower than that for randomly simulated puncta (Figure 2G), suggesting that autophagosomes are delivered to and degraded in tubules of the tAL network.

Altogether, we identified a distinct and highly tubulated autolysosomal compartment that expands with muscle remodeling. This tubulated autolysosome is uniquely marked by Stx17 and exhibits degradative capacity throughout the tubular network. Going forward, we refer to this structure as the tubular <u>a</u>utolysosomal (tAL) network.

### Formation of the tubular autolysosomal network requires lysosomal function, independent of mTOR activity

To test whether the autolysosomal degradative function is required for formation of the tubulated network, we examined effects from knockdown of lysosomal functions, *Spinster, TRPML* and *Vha68-3* (Dermaut et al., 2005; Wong et al., 2012; Mauvezin et al., 2015). Strikingly, knockdown of each disrupted the GFP:Stx17 tubular network (Figure 2H–I), suggesting that lysosomal homeostasis and/or cargo degradation is critical for network tubulation. Since uptake of extracellular DQ Red-BSA must occur for eventual colocalization at autolysosomes (Figure 2E), we postulated that formation of the tAL network may also depend on endocytic delivery to lysosomes (Guha et al., 2003). To test this possibility, we conditionally disrupted *shibire*, the sole fly ortholog of dynamin involved in endocytic uptake for a significant portion of cell surface cargos delivered to lysosomes (Kosaka and Ikeda, 1983). Flies with the temperature-sensitive mutation, *shi*^*ts1*^, were reared at permissive temperature (19°C) until 12 h APF, shifted to restrictive temperature (29°C), and then examined at 20 h APF (Figure 2—figure supplement 2A). A block in *shibire* function scarcely affected Spin:RFP organization (Figure 2—figure supplement 2B–C), suggesting that dynamin-dependent endocytosis does not substantially contribute to formation of the tAL network.

In the process of ALR, autolysosomal tubulation depends on reactivation of mTORC1 activity in response to efflux of autophagic degradation products (Yu et al., 2010). To test if mTOR activity is involved in the formation of the tAL network, mTOR activity was forcibly inactivated or activated in DIOMs (Dibble and Cantley, 2015). Inactivation of mTOR by *Tor* or *Rheb* RNAi resulted in thinner DIOMs (Figure 2— figure supplement 3A–C), however, the formation of the tAL network was largely unaffected at both 20 h and 4 d APF (Figure 2—figure supplement 3C–D). In mammalian cells, forced activation of mTOR activity could suppresse the loss of ALR tubulation due to *Spinster* RNAi (Rong et al., 2011). Activation of mTOR by *Tsc1* RNAi or *Rheb* overexpression led to thicker DIOMs (Figure 2—figure supplement 3E–F), however, was unable to suppress loss of the tAL network in *Spin* RNAi conditions (Figure 2—figure supplement 3G–H). Thus, our data suggest that mTOR activity is not essential for the formation of the tubulated autolysosome network in muscle.

### Formation of the tubular autolysosomal network depends on autophagy, but not the Atg12 conjugation system

Next, we asked if autophagy is required for formation of the tAL network. We tested the requirements for at least one *ATG* gene from each of the six functional protein units involved in autophagy (Mizushima et al., 2011), as well as genes required for the fusion between autophagosomes and lysosomes (Lőrincz and Juhász, 2019). RNAi of *Atg1, FIP200, Atg9, Atg18, Vps34, Stx17, SNAP29*, or *Vps39* each severely blocked formation of the tubular autolysosomal network (Figure 3A–B). In contrast, *Atg5, Atg7*, or *Atg12* RNAi showed only a minimal effect on the tubular network. We obtained similar results for the genes tested using tAL compartment markers, Spin:RFP (Figure 3A–B) or mCherry:Stx17 (Figure 3—figure supplement 1A–B). These results suggest the importance of autophagy, surprisingly without the Atg12 conjugation system, for tAL network formation.

**Figure 3.**
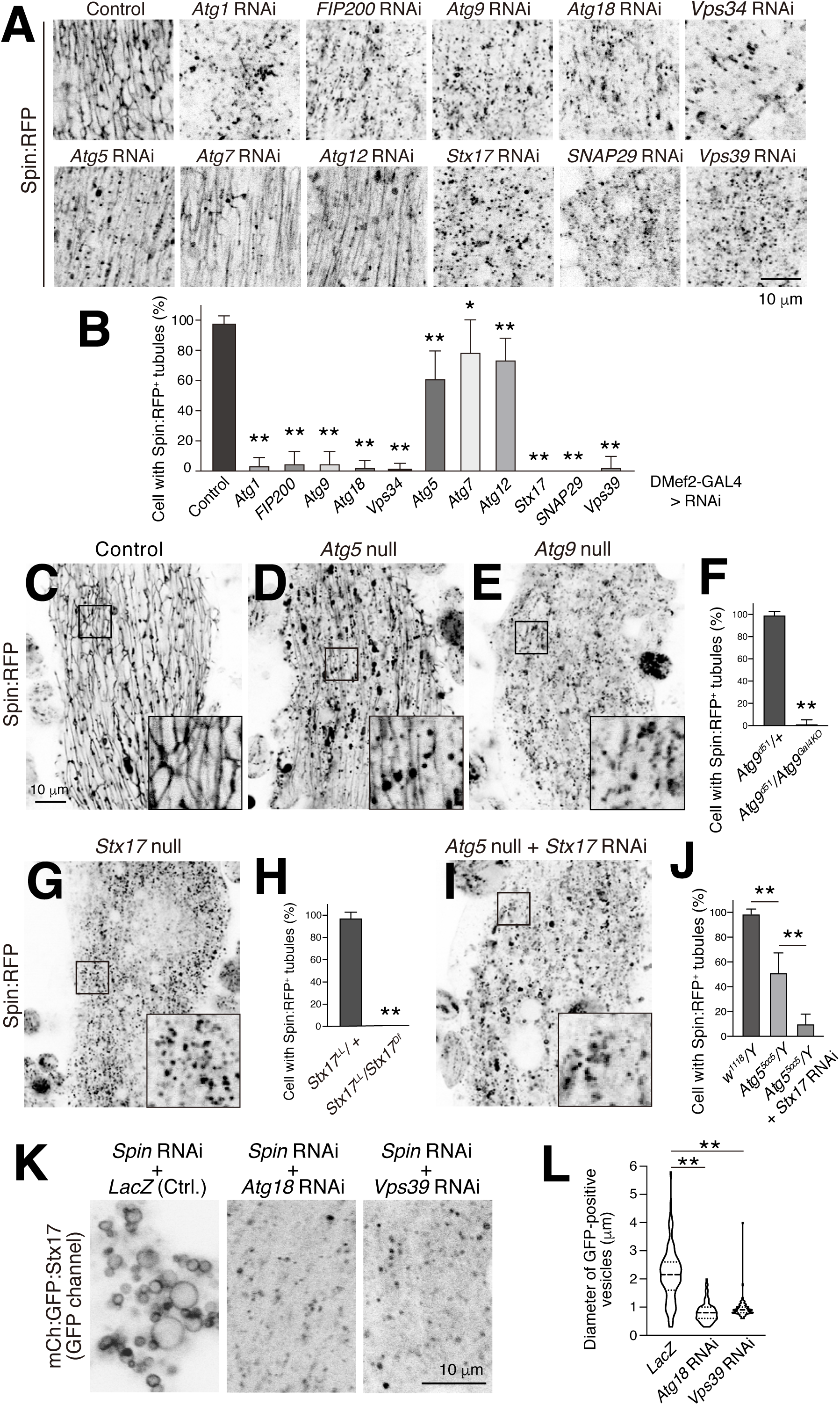
Formation of the tubular autolysosomal network depends on autophagy, but not the Atg12 conjugation system. (A and B) Effect of RNAi of autophagy-related (*ATG*) genes on Spin:RFP-positive tubular network in 20 h APF DIOMs (A). Mean percent + SD of DIOMs with more than 5 μm Spin:RFP-positive tubules, n=10. *, p<0.05; **, p<0.001 (Dunnett’s test) (B). (C-E) Spin:RFP-positive tubules in control (C), *Atg5* null (D), or *Atg9* null (E) DIOMs at 20 h APF. (F) Mean percent + SD of DIOMs with Spin:RFP-positive tubules in control or *Atg9* null mutant, n=10. **, p <0.001 (Student’s *t*-test). (G and H) Spin:RFP-positive tubules in *Stx17* null DIOMs at 20 h APF (G). Mean percent + SD of DIOMs with Spin:RFP-positive tubules in control or *Stx17* null, n=10. **, p <0.001 (Student’s *t*-test) (H). (I and J) Spin:RFP in *Stx17*-knockdowned *Atg5* null DIOMs at 20 h APF (I). Mean percent + SD of DIOMs with Spin:RFP-positive tubules in control, *Atg5* null, or combination of *Atg5* null and *Stx17* RNAi, n=10. **, p<0.001 (Tukey’s test) (J). (K and L) Co-RNAi of *Spin* and *Atg18* or *Vps39* on mCherry:GFP:Stx17 in 20 h APF DIOMs (K). Violin plot of the diameter of mCherry:GFP:Stx17-positive vesicles in each genotype, n=100. **, p<0.001 (Dunnett’s test) (L).

To verify that the categories of *ATG* gene results were not simply due to variability with hypomorphic RNAi conditions, we examined null mutants for *Atg5, Atg9*, and *Stx17* (Kim et al., 2016; Wen et al., 2017; Takáts et al., 2013). Consistent with the RNAi results, loss of *Atg9* or *Stx17* functions fully blocked tAL formation, while *Atg5* null mutants only partially reduced the extent of the tubulated network (Figure 3C–H and Figure 3—figure supplement 1C–E). The tubulated autolysosome still present in *Atg5* null mutant DIOMs was dependent on *Stx17* function (Figure 3I–J). The size of the Stx17 vacuoles seen upon *Spinster* RNAi was significantly reduced by the combined knockdown of autophagy functions, *Atg18* or *Vps39* (Figure 3K–L), suggesting that the Stx17 compartment size depends on membrane flux through autophagy. From these results, we conclude that autophagy – independent of the Atg12 conjugation system – is necessary for formation of the tAL network.

A tubular lysosomal network has been reported in larval body wall muscles (Johnson et al., 2015). The authors reported that autophagy is not a prerequisite for the tubulation, since Atg7 RNAi did not affect it. Atg7 is an E1 enzyme for both the Atg12 and Atg8 ubiquitin-like conjugation systems (Juhász et al., 2007). As in larval muscles, Atg7 also was not required for formation of the tAL network in DIOMs (Figure 3A–B). Thus, we predicted that the tubular lysosome in larval body wall muscles may also depend on core *ATG* genes, but not Atg5, Atg7 or Atg12. As reported, Spin:RFP-positive tubular lysosomes were observed close to the muscle cell surface in control and *Atg7* RNAi larval body wall muscles (Figure 3—figure supplement 2). In contrast, the tubular network was disrupted by *Atg1, Atg18* or *Stx17* RNAi (Figure 3—figure supplement 2), indicating that autophagy is also essential for the tubular lysosomal network in larval body wall muscles.

### Ultrastructure supports that autophagosome is membrane source for tAL network

To analyze the ultrastructure of the tAL network, DIOMs were cut longitudinally and examined by transmission electron microscopy (TEM). Lysosomes and autolysosomes appear as electron-dense structures by TEM. Consistent with this, we observed electron-dense tubular structures in both control (Figure 4A, 4E-F) and *Atg5* null mutant DIOMs (Figure 4D). The diameter of the tubules seen by TEM were ranged between approximately 50-100 nm. In contrast, mostly short or spherical electron-dense structures were observed in *Atg9* or *FIP200* null (Kim et al., 2013) (Figure 4B–C). On the other hand, large vacuolated structures accumulated upon *Spin* or *TRPML* RNAi (Figure 4G–H), consistent with the light microscopy results (Figure 2H). Autophagosome-like double-membrane vesicles were observed in *Atg5* null DIOMs (Figure 4—figure supplement 1), as similarly reported in *Atg3* KO mammalian cells (Tsuboyama et al., 2016), indicating that *Atg5* is not essential for elongation of the autophagic membrane in DIOMs at 1 d APF. Since autophagosome-like double-membrane vesicles and the tAL network are still formed in *Atg5* mutant muscles, yet the tAL network depends on Stx17 (Figure 3I–J), we conclude that autophagic membrane is a primary membrane source for tAL network formation in both wildtype and *Atg5* null muscle cells.

**Figure 4.**
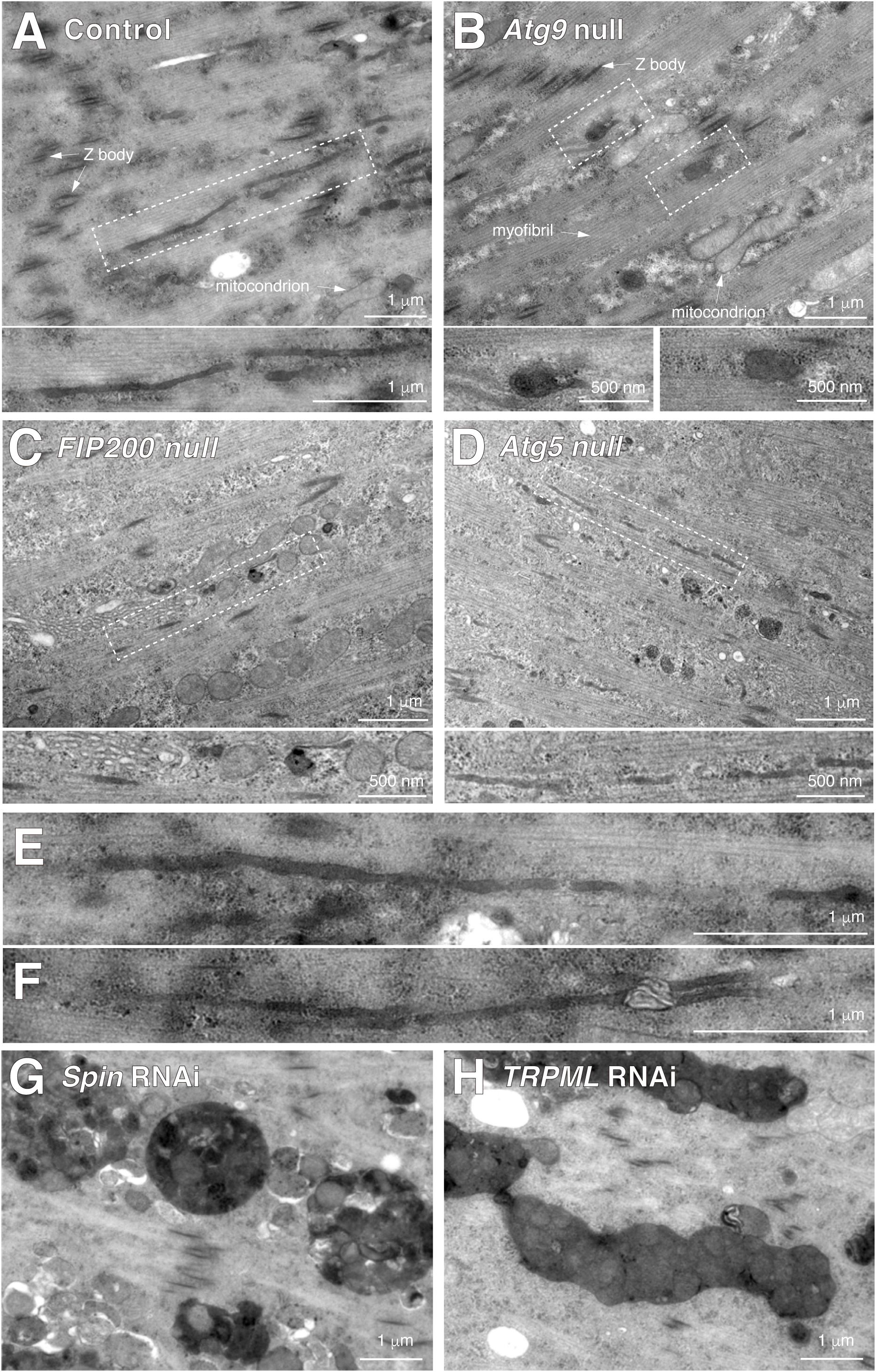
Ultrastructure supports that autophagosome is membrane source for tAL network. TEM images of 20 h APF DIOMs of control (A, E, F), *Atg9* null (B), *FIP200* null (C), *Atg5* null mutant (D), *Spin* RNAi (G), or *TRPML* RNAi (H). Typical examples of electron-dense membranous tubular structures in control DIOMs (E and F).

### Extent of tubular network correlates with muscle remodeling ability

As shown above, there was a significant difference in the efficiency of tAL network formation between disruption of genes core to autophagy versus genes in the Atg12 conjugation system (Figures 3–4). We next compared whether the same loss-of-function conditions also differentially impacted muscle remodeling. Knockdown of *FIP200, Atg9*, or *Atg18* had a noticeable effect on DIOM shape at 4 d APF, after remodeling is completed. However, knockdown of *Atg5* or *Atg12* did not affect muscle shape (Figure 5A–B), demonstrating again two distinct categories of *ATG* phenotypes with DIOM remodeling.

**Figure 5.**
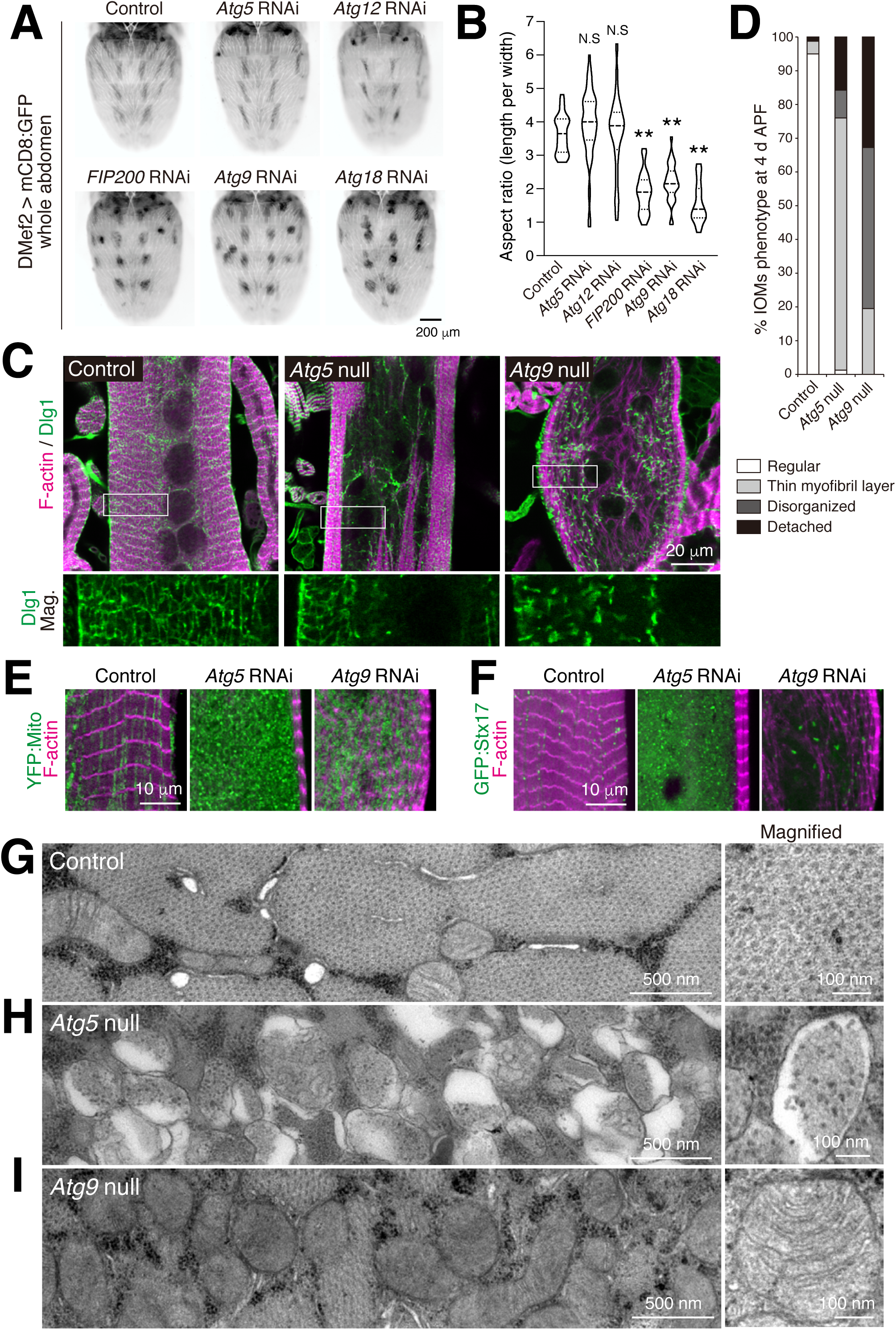
Extent of tubular network correlates with muscle remodeling ability. (A and B) Effect of RNAi of *ATG* genes on the shape of DIOMs at 4 d APF (A). Violin plot of the aspect ratio of DIOMs, n>50. N.S, not significant; **, p<0.001 (Dunnett’s test) (B). (C and D) T-tubule (Dlg1, green) and myofibril (F-actin, magenta) organization in DIOMs of control, *Atg5* null, or *Atg9* null (C). Quantification of DIOM phenotypes in control, *Atg5* null, or *Atg9* null DIOMs, n>10. (D). Regular, straight DIOM with organized myofibrils and T-tubules; Thin myofibril layer, straight DIOM with thin myofibrils; Disorganized, irregular shaped DIOM with disorganized myofibrils and T-tubules; Detached, detached and rounded DIOM. (E) Mitochondria (YFP:Mito, green) and myofibrils (F-actin, magenta) in 4 d APF DIOMs of control, *Atg5*, or *Atg9* RNAi. (F) Autophagic membranes (GFP:Stx17, green) and myofibrils (F-actin, magenta) in 4 d APF DIOMs of control, *Atg5*, or *Atg9* RNAi. (G-I) TEM images of 4 d APF DIOM transverse-sections of control (G), *Atg5* null (H), or *Atg9* null (I).

We characterized each of the two phenotypic subgroups further at the organelle level using *Atg5* and *Atg9* mutant conditions. Following completion of remodeling, control DIOMs had well-organized myofibrils (F-actin) and T-tubules (Dlg1) at 4 d APF. While the shape was nearly normal for the *Atg5* null DIOMs, they contained both an organized peripheral layer of myofibrils and T-tubules and a disorganized central region at 4 d APF (Figure 5C). In contrast, *Atg9* null animals had irregularly shaped DIOMs with more extensively disorganized myofibrils and fragmented Dlg1-positive structures throughout the cells. Mitochondria accumulated with either *Atg5* or *Atg9* RNAi conditions (Figure 5E), suggesting a block in mitophagy for both conditions. GFP:Stx17-positive autophagic structures, however, only accumulated in *Atg5* RNAi but not in *Atg9* RNAi muscles (Figure 5F).

We performed TEM of transverse sections through remodeled DIOMs (Figure 5—figure supplement 1A). Control DIOMs were filled with myofibrils and organized organelles recognizable by ultrastructure, such as mitochondria and T-tubules (Figure 5G and Figure 5—figure supplement 1B). The *Atg5* null DIOMs instead were filled with thousands of autophagic membranes (Figure 5H and Figure 5—figure supplement 1C and 1E), likely representative of the numerous GFP:Stx17-positive vesicles also shown to accumulate (Figure 5F). In contrast, *Atg9* null DIOMs were filled with mitochondria but lacked any recognizable autophagic compartments (Figure 5I and Figure 5—figure supplement 1D), similar to phenotypes previously described for *Atg1* or *Atg18* RNAi (Fujita et al., 2017). Collectively, these data demonstrate that there are two distinct loss of *ATG* phenotypes in DIOM remodeling, with the efficiency of tAL network formation correlating with the phenotypic severity of muscle remodeling: only a slightly reduced tAL network correlating with partially organized muscle (*Atg5* mutant), and a fully disrupted tAL network associated with more completely disorganized muscle (*Atg9* mutant).

### The tubulated lumen is continuous to synchronize autolysosomal capacity with muscle remodeling

What is an advantage of having autolysosome organization into a tubular network over numerous isolated spherical vesicles? The interconnected tAL network could enable synchronous degradative activity across broad regions of the relatively large muscle cells. To investigate the continuity of the tAL network, we performed fluorescence recovery after photobleaching (FRAP) analysis of fly cathepsin L, Cp1. At 24 h APF, GFP:Stx17 marked large, rounded intersections between several tubule branches that were filled with Cp1:mKO (Figure 6A–B). In control DIOMs, Cp1:mKO signal at the intersecting branchpoints recovered over several minutes after bleaching (Figure 6C and 6E). In contrast, the signal of Cp1:mKO was not recovered in discontinuous vacuoles in *Spin* RNAi DIOMs (Figure 6D–E). These results show that the lumen of the tAL network is continuous and allows protein contents between tubules to intermix.

**Figure 6.**
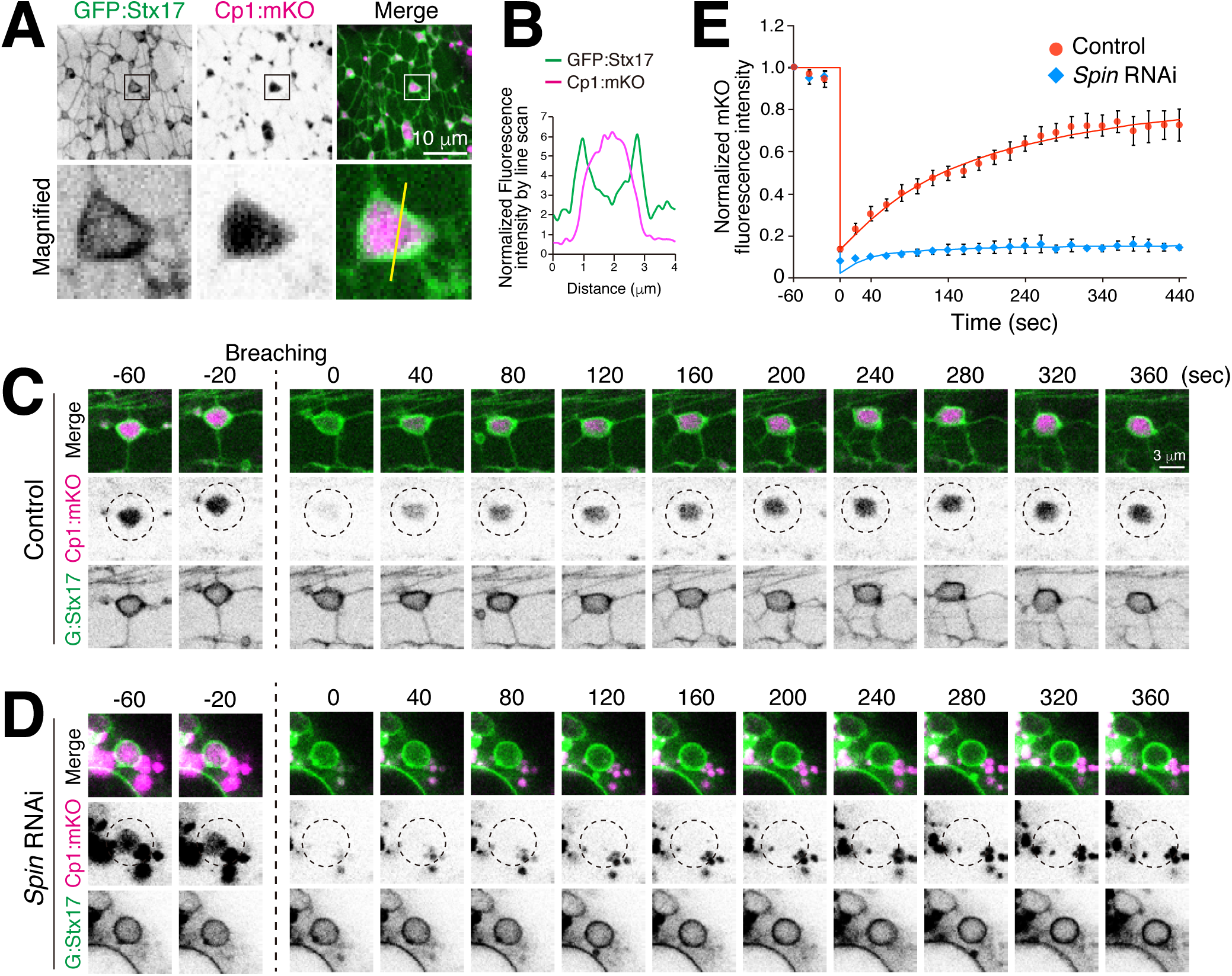
The tubulated autolysosome lumen is continuous and intermixed. (A and B) Colocalization of GFP:Stx17 and Cp1:mKO in 24 h APF DIOM (A). Line plot profiles of the yellow line in panel A (B). (C and D) FRAP analysis of Cp1:mKO in 24 h APF DIOMs of control (C) or *Spin* RNAi (D). (E) Quantification of panels C and D. The average ± SD is shown, n=5.

We next tested whether lysosomal activity is more homogeneous across a tAL network than that found amongst multiple individual lysosomes over a similar muscle area. LysoTracker Red or DQ Red-BSA was injected into pupae expressing GFP:Stx17 to stain acidified compartments. Both indicators stained small discontinuous vesicles in 12 h APF DIOMs (Figure 7A and 7C) and the intersections of the tAL network in 24 h APF DIOMs (Figure 7A and 7C), respectively. The acquired confocal images were binarized, extracted objects, and the mean intensities of each LysoTracker Red or DQ Red-BSA-positive object were measured. As predicted, the intensities of LysoTracker Red or DQ-BSA were more heterogeneous in discontinuous lysosomes at 12 h APF and more homogeneous in the tAL network at 24 h APF (Figure 7B and 7D). These results indicate that the tAL network synchronizes the degradative compartments.

**Figure 7.**
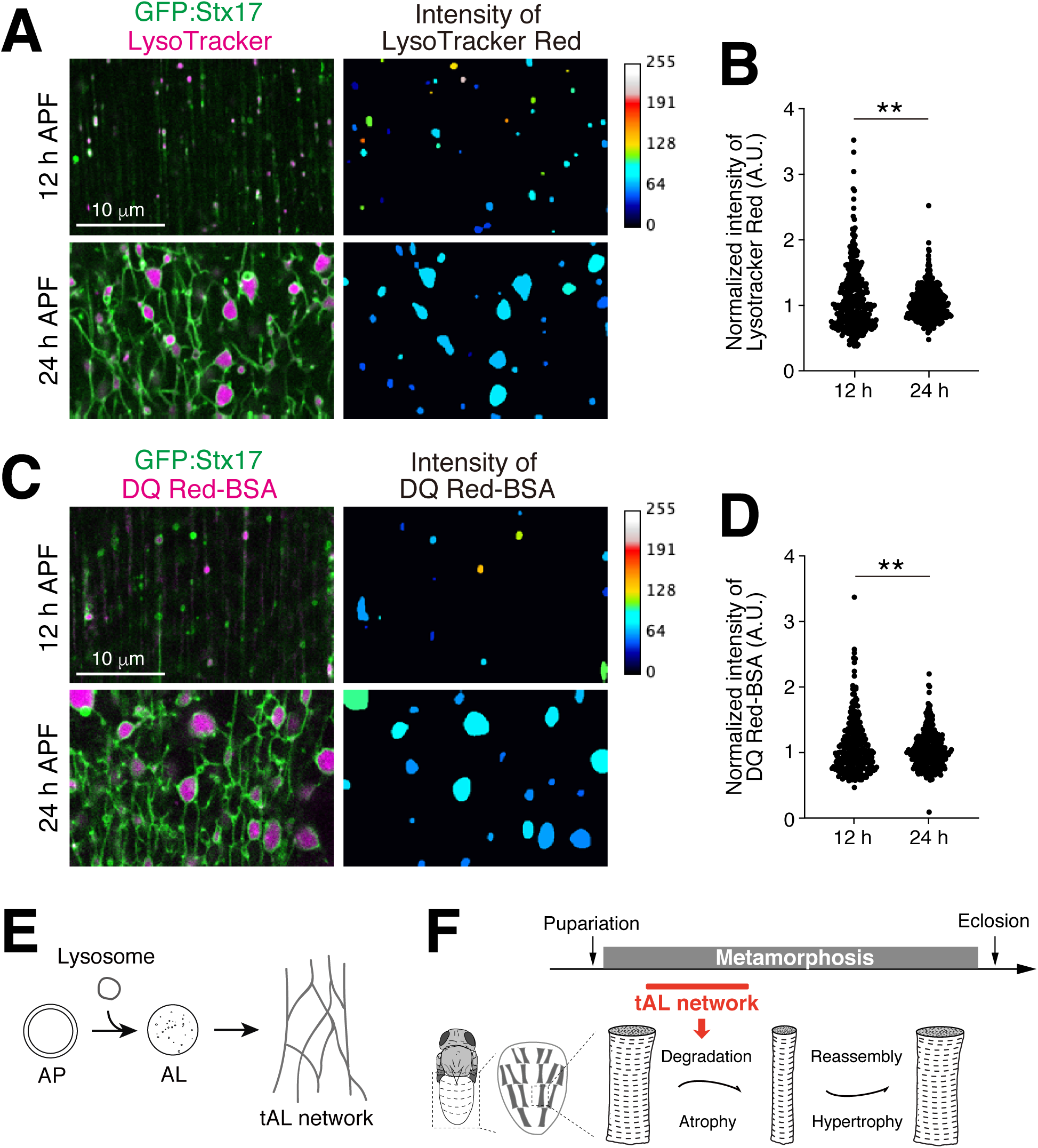
The tubular autolysosomal network synchronizes the degradative compartments. (A and B) Micro-injected LysoTracker Red and GFP:Stx17 in 12 or 24 h APF DIOMs. The intensity map shows a representative image of the intensities of LysoTracker Red-positive objects. The median intensity in each image was set as 60 (A). The intensities of LysoTracker-positive objects were shown in a scatter plot, n>250. The median intensity was set as one for each image. **, p<0.001 (*F*-test) (B). (C and D) Micro-injected DQ Red-BSA and GFP:Stx17 in 12 or 24 h APF DIOMs. The intensity map shows the intensities of DQ Red-BSA-positive objects. The median intensity in each image was set as 60 (C). The intensities of DQ Red-BSA-positive objects were shown in a scatter plot, n>250. The median intensity was set as one for each image. **, p<0.001 (*F*-test) (D). (E) Both autophagy and lysosomal degradative activity are required for the tAL network formation. (F) The tAL network functions in the degradation of organelles in the remodeling of Drosophila abdominal muscles.

## Discussion

Tubular lysosomes have been known since over 40 years ago, but the mechanisms of how they are formed and their significance are mostly unknown. Here, we found expansion of a tubular autolysosomal network in the Drosophila abdominal muscles during metamorphosis. The tubular network appears transiently and is fully functional as a degradative organelle. The formation of the network depends on not only autophagy but also the degradation capacity of acidic organelles. FRAP analysis of cathepsin L revealed that the lumen of the tubular network is continuous and intermixed. As far as we know, the autophagy-dependent activity of a tubular lysosomal network has not been reported. Accordingly, we designate the structure as the tubular autolysosomal network, or tAL network.

In contrast to the already known tubular lysosomes, the tAL network is an autophagy-related organelle (Figure 7E). First, we found the tAL network is uniquely marked with Stx17, previously known as an autophagosomal SNARE (Figure 1). However, the tAL network has characteristics of autolysosomes and not autophagosomes (Figure 2). Further, the formation of the tAL network depends on autophagy, and the loss of a series of *ATG* genes led to the accumulation of only small spherical lysosomes (Figures 3–4, Figure 3—figure supplement 1, and, Figure 7— figure supplement 1A). To our surprise, components of Atg12 and Atg8 systems were not essential for formation of the tAL network (Figures 3–4 and Figure 7—figure supplement 1B).

How is the tAL network formed in absence of *Atg5*, a component of the Atg12 system? The formation of the tAL network in wildtype and *Atg5* deficient muscles both depended on *Stx17* (Figure 3I–J). Further, elongated autophagic membranes accumulated in the *Atg5* null DIOMs (Figure 4—figure supplement 1). We propose that the fusion between autophagic membranes and lysosomes results in the formation of the tAL network (Figure 7—figure supplement 1B). We predict that the autophagic membrane but not the endocytic membrane is the dominant membrane source, since inhibition of the fusion of autophagosome and lysosome wholly prevented tAL network formation (Figure 3). Consistent with this notion, a block in dynamin-dependent endocytosis hardly affected the tAL network (Figure 2—figure supplement 2A-C). The formation of a tAL network in *Atg5* null DIOMs could be explained by diminished yet ongoing leaky autophagic flux. Because almost all autophagic flux assays in Drosophila utilize Atg8, lipidation of which depends on Atg5, we could not judge the autophagic flux in the *Atg5* null mutant; however, the *Atg5* null mutant might degrade autophagic cargoes to some extent. Our TEM analysis revealed the existence of elongated autophagic membrane in the *Atg5* null DIOMs (Figure 4—figure supplement 1). Even if the autophagosomal membranes are incompletely closed in the *Atg5* null condition, which would block autophagy, membrane fusion with multiple lysosomes could result in autophagosomal membrane closure. It is critical to establish an Atg8-independent autophagic flux assay to test the hypothesis.

It has been reported that Drosophila larval muscles have a tubular lysosomal network (Johnson et al., 2015). Since *Atg7* RNAi did not block the tubulation, it was thought that autophagy is dispensable for the tubular network formation. Through the analysis of the tAL network in pupal DIOMs, we found that *Atg7* and other components of two ubiquitin-like conjugation systems were dispensable for formation of the tAL network, the same as the tubular lysosomal network in larval body wall muscle (Figures 3–4). Further, we revealed that the tubular lysosomal network in larval muscles also depended on *Atg1, Atg18*, and *Stx17* (Figure 3—figure supplement 2), in agreement with our result in pupal muscle. Thus, it is reasonable to think that the tAL network and the tubular lysosome in larval muscles are closely related structures. Although we first discovered the tAL network in DIOMs during metamorphosis, we observed a similar structure to varying degrees in other muscles, such as the adult indirect flight muscles that form in the pupal thorax and dorsal longitudinal muscles in the abdomen.

The tAL network has different characteristics from ALR tubules, which are proto-lysosomes and do not have degradation capacities (Chen and Yu, 2013). The tAL exhibits acidification and degradative function throughout the tubular network, and both endocytosed DQ Red-BSA and autophagosomes were degraded in the structure (Figure 2). Although the tAL network and the ALR tubules have these dissimilarities, both are autolysosome-related compartments. More work is needed to address whether the formation of the tAL network also depends on factors involved in ALR, such as clathrin, PI(4,5)P_2_, microtubules, and Kinesin 1 (Rong et al., 2012; Du et al., 2016).

To our surprise, Stx17 localizes to the autolysosomal compartment in the remodeling muscle (Figure 2). So far, it is known that Stx17 localizes to the autophagosome and detaches just after fusion with the lysosome in mammalian cells (Tsuboyama et al., 2016). Stx17 is a unique SNARE protein, which has a hairpin-like two *α*-helixes in its carboxy-terminus. Through the acidic *α*-helixes, Stx17 is specifically recruited to the autophagic membranes (Itakura et al., 2012; Takáts et al., 2013). Similar to Stx17, the ER morphogens, such as Reticulons and REEPs, have hairpin-like membrane-anchoring domains, which are inserted into and drive deformation of the ER membrane like wedges (Park and Blackstone, 2010). Therefore, it is possible to speculate that Stx17 induces the tubulation of membranes, like the ER morphogens. Further studies are required to reveal the mechanisms shaping positive membrane curvature and a highly branched network of the tAL network.

What is the advantage to forming a tubular autolysosomal network? Our data suggests that there are two merits of the tAL network over spherical autolysosomes. First, is the expansion of the surface area. TEM data indicates that the diameter of the tubes is ranged between 50-100 nm (Figure 4). If we compare a ratio of surface area per volume of a 70-nm-diameter tube and 500-nm-diameter spherical vesicle, the tube has ∼5 times higher score than that of the vesicle (Figure 7—figure supplement 1C). The formation of a tAL network would increase the chance of docking and fusion of autophagosomes with the degradative compartments. Autophagy is massively induced with the onset of DIOM remodeling (Fujita et al., 2017), and autophagosomes fuse with and are degraded in the tAL network (Figure 2F–G). Thus, the tAL network would allow handling the extremely high autophagic flux in the relatively large, multinucleated muscles cells. We propose that there is positive feedback on the autophagic degradation in the remodeling DIOMs. The induction of autophagy to higher levels leads to the tAL network formation, which is able to more efficiently fuse with and degrade more autophagosomes than are spherical lysosomes. Alternatively, the expanded surface by tubulation may be advantageous for the process of microautophagy, in which the lysosomes directly engulf cytosolic materials by membrane invagination (Oku and Sakai, 2018).

The second benefit to tubulated autolysosomes is the synchronization of the degradative compartments in the cell. FRAP analysis of Cp1:mKO revealed that the lumen of the tAL network is continuous and intermixed (Figure 6C–E). With DIOM remodeling, organelles are disassembled in the early pupal stage and then reassembled in the late pupal stage (Figure 7F). For the regulated muscle remodeling, all events, such as degradation and signaling from the degradative organelle, must be synchronized.

Because muscles are massive cells, it is difficult to synchronize a number of spherical lysosomes over a wide cellular region. The formation of the tubular network would synchronize the degradative compartments in the cell (Figure 7—figure supplement 1C). Consistent with this notion, we showed that the tAL network activity is more homogeneous than the activity of discontinuous, spherical lysosomes (Figure 7A-D).

Both Atg5 and Atg9 seem to be pivotal for autophagy in DIOMs; however, the null mutants showed distinct phenotypes on both the tAL network and muscle remodeling (Figure 5). In contrast to Atg9, Atg5 was required but not essential for the formation of the tAL network (Figures 3–4). Again, *Atg5* null induced a milder phenotype than *Atg9* null on the muscle remodeling (Figure 5). Further, we observed that the muscle remodeling was also severely affected in *Spin* or *Vha68-3* RNAi, which induced loss of the tAL network in 1 d APF DIOMs (Figure 2H). These correlations suggest that the extent of the tAL network plays an important role in DIOM remodeling (Figure 7F).

The existence of the tubular lysosome has been known in a variety of tissues; however, the mechanism and biological significance of the tubulation remain obscure and highly speculative. It is likely that mammalian muscles also have the tubulated lysosomes (Robinson et al., 1986; Okada et al., 1986). Because the tubular lysosomes, including the tAL network in DIOMs, are fragile and highly sensitive to dissection and fixation processes, live imaging is an essential technique for the analysis of the structure. Since fly abdominal muscles are located close to the transparent overlying cuticle, we succeeded in observing the tubular autolysosome in live animals through the cuticle. Hence, Drosophila is an ideal model system to analyze the tubular lysosomal network and the dynamics of muscle remodeling. Our findings in this study provide new insights into the mechanisms of the morphogenesis of lysosomes as well as regulation of fundamental membrane trafficking pathways, such as autophagy and endocytosis. We predict that the expansion of surface area and synchronization are the keys to understanding the tubular lysosomes. Identification of genes that are specifically required for the tubular network would be the next crucial step and answer the fundamental question, why the lysosomes dynamically change shape in certain conditions.

## Materials and methods

### Reagents and antibodies

The following reagents were used: Alexa Fluor 546 Phalloidin conjugate (1.0 U/mL; Invitrogen), LysoTracker Red DND-99 (Thermo Fisher Scientific, Waltham, MA), and DQ-Red BSA (Thermo Fisher Scientific, Waltham, MA). The following antibodies were used: mouse anti-fly Dlg1 (1:200; clone 4F3; Developmental Studies Hybridoma Bank, Iowa City, IA) and anti-mouse IgG Alexa Fluor 488 conjugate (1:1000; Thermo Fisher Scientific, Waltham, MA).

### Drosophila strains

Flies were reared at 25°C, unless stated. For muscle-targeted gene expression, DMef2-GAL4 driver was used. *UAS-LacZ* was used as a control in RNAi experiments. Genotypes in figures were described in supplemental table 1. All genetic combinations were generated by standard crosses. Genotypes of flies used in this study include the following: (1) *Atg5*^*5cc5*^/*FM7 actin-GFP* (from JH. Lee; *Atg5* null), (2) *Atg9*^*Gal4KO*^/*CyO-GFP* (from G. C. Chen; *Atg9* null), (3) *Atg9*^*d51*^/*CyO-GFP* (from G. C. Chen; *Atg9* null), (4) *w; FIP200*^*3F5*^/*TM6B-GFP* (from J.H. Lee; *FIP200* null), (5) *w; FIP200*^*4G7*^/*TM6B-GFP* (from J.H. Lee; *FIP200* null), (6) *w; UASp-mCherry:GFP:Atg8a* (from H. Stenmark), (7) *y w; UAS-GFP:Atg8a* (from T. Neufeld), (8) *y w; UAS-mCherry:Atg8a* (from T. Neufeld), (9) *w; UAS-Spinster:myc:RFP*/*CyO* (From G. Davis), (10) *w; UAS-IR-Atg1* (from G. C. Chen), (11) *w, shi*[*1*] (Bloomington Drosophila Stock Center, BDSC 7968; *shibire* temperature-sensitive allele), (12) *w; Df*(*3L*)*Exel8098*/*TM6B, Tb*[*1*] (BDSC 7922; Stx17 deficiency), (13) *y w; P*{*w*[*+mC*]*=GAL4-Mef2.R*}*3* (BDSC 27390), (14) *w; P*{*w*[*+mC*]*=UAS-lacZ.B*}*melt*[*Bg4-1-2*] (BDSC 1776), (15) *y w; P*{*y*[*+t\**] *w*[*+mC*]*=UAS-Lifeact-Ruby*}*VIE-19A* (BDSC 35545), (16) *w; P*{*w*[*+mC*]*=UAS-mCD8::GFP.L*}*LL5, P*{*UAS-mCD8::GFP.L*}*2* (BDSC 5137), (17) *w; P*{*w*[*+mC*]*=UAS-Rheb.Pa*}*3* (BDSC 9689), (18) *y v; P*{*y*[*+t7.7*] *v*[*+t1.8*]*=TRiP.HMS02818*}*attP40* (TRiP, BDSC 44098; TRPML RNAi), (19) *y v; P*{*y*[*+t7.7*] *v*[*+t1.8*]*=TRiP.HMS01611*}*attP2*/*TM3, Sb*[*1*] (TRiP, BDSC 36918; FIP200 RNAi), (20) *y v; P*{*y*[*+t7.7*] *v*[*+t1.8*]*=TRiP.HMS01246*}*attP2* (TRiP, BDSC 34901; Atg9 RNAi), (21) *y v; P*{*y*[*+t7.7*] *v*[*+t1.8*]*=TRiP.HMS01358*}*attP2*/*TM3, Sb*[*1*] (TRiP, BDSC 34369; Atg7 RNAi), (22) *y v; P*{*y*[*+t7.7*] *v*[*+t1.8*]*=TRiP.HMS01153*}*attP2* (TRiP, BDSC 34675; Atg12 RNAi), (23) *y v; P*{*y*[*+t7.7*] *v*[*+t1.8*]*=TRiP.GL00012}attP2* (TRiP, BDSC 35144; Tsc1 RNAi), (24) *y v; P*{*y*[*+t7.7*] *v*[+*t1.8*]*=TRiP.HMS00923*}*attP2* (TRiP, BDSC 33966; Rheb RNAi), (25) *y v; P*{*y*[*+t7.7*] *v*[*+t1.8*]*=TRiP.GL00156*}*attP2* (TRiP, BDSC 35578; Tor RNAi), (26) *y v; P*{*y*[*+t7.7*] *v*[*+t1.8*]*=TRiP.JF01937*}*attP2* (TRiP, BDSC 25896; Stx17 RNAi), (27) *y v; P*{*y*[*+t7.7*] *v*[*+t1.8*]*=TRiP.JF01883*}*attP2* (TRiP, BDSC 25862; SNAP29 RNAi), (28) *y v; P*{*y*[*+t7.7*] *v*[*+t1.8*]*=TRiP.HMS02438*} (TRiP, BDSC 42605; Vps39 RNAi), (29) *Sco*/*CyO; P*{*UAS-Cp1.mKO2*}*3* (VDRC 309010), (30) *P*{*Mef2-GAL4*}*3, P*{*UAS-Cp1.mKO2*}*3*/ *TM6B, Tb*[*1*] (VDRC 309005), (31) *w; UAS-IR-Atg18*^*KK100064*^ (VDRC 105366; Atg18 RNAi), (32) *w; UAS-IR-Spinster* (NIG-Fly 8428R-4; Spin RNAi), (33) *w; UAS-IR-Vha68-3* (NIG-Fly 5075R-1; Vha68-3 RNAi), (34) *w; UAS-IR-Vps34* (NIG-Fly 5373R-2; Vps34 RNAi), (35) *w; UAS-IR-Atg5* (NIG-Fly 1643R-2; Atg5 RNAi), (36) *w; UAS-mCD8:GFP; DMef2-GAL4, UAS-Dcr2*, (37) *w; DMef2-GAL4, UAS-Dcr2, w; UAS-Dcr2; DMef2-GAL4, UAS-GFP:Stx17*^*4*^, and (39) *w; UASt-GFP:Stx17*^*4*^. New genotypes generated during this study include the following: (40) *w; DMef2-GAL4, UAS-GFP:Stx17*^*4*^, (41) *w; UAS-Spin:myc:RFP*/*CyO; DMef2-GAL4, UAS-Dcr2*/*TM6C Sb Tb*, (42) *w; Atg9*^*d51*^, *UAS-Spin:myc:RFP*/*CyO; DMef2-GAL4, UAS-Dcr2*/*TM6C Sb Tb*, (43) *w; UAS-Spin:myc:RFP*/*CyO; DMef2-GAL4, Stx17*^*LL06330*^/*TM6C Sb Tb*, (44) *w; UAS-Spin:myc:RFP*/*CyO; DMef2-GAL4, UAS-IR-Stx17*^*TRiP.JF01937*^/*TM6B Hu Tb*, (45) *w; UAS-IR-Spinster* ^*NIG.8428R-4*^, *UAS-mCherry:GFP:Stx17*/*CyO; DMef2-GAL4, UAS-Dcr2, tub-GAL80*^*ts*^, (46) *w; UAS-Dcr2; DMef2-GAL4, sqh-YFP:Mito*^*BDSC7194*^/*TM6C Sb Tb*, (47) *w; UAS-Dcr2*/*CyO; DMef2-GAL4, UAS-Cp1:mKO, UAS-GFP:Stx17*, (48) *w; UAS-Dcr2; DMef2-GAL4, UAS-mCherry:Stx17*/*TM6C Sb Tb*, (49) *w; Sp*/*CyO; DMef2-GAL4, UAS-mCherry:Stx17*/*TM6C Sb Tb*, (50) *w; Atg9*^*d51*^/*CyO-GFP; DMef2-GAL4, UAS-mCherry:Stx17*/*TM6C Sb Tb*, and (51) *w; Atg9*^*d51*^/*CyO-GFP; DMef2-GAL4, UAS-mCherry:Stx17*/*TM6C Sb Tb.*

### DNA engineering

Standard molecular biology techniques were used to construct plasmid vectors. For multiple-fragment in vitro assembly, pCM43b vector was digested by *Eco*RI and *Not*I. mCherry and GFP:Stx17 were amplified using following primer sets; 5’-AGGGAATTGGGAATTCACCATGGTTTCAAAAGGTGAAG-3’ and 5’-GCTTCCTCCTCCTCCCTTGTACAGCTCGTCCATGCCGCC-3’ for mCherry, 5’-GGAGGAGGAGGAAGCATGGTGAGCAAGGGCGAG-3’ and 5’-TCCTCTAGTGCGGCCTCATTCTGGCTTCTCTTTTAGC-3’ for GFP:Stx17. The three DNA fragments were assembled using In-fusion HD cloning kit (Takara, Kusatsu, Japan). The resultant DNA construct (pCM43b-mCherry:GFP:Stx17) was validated by sequencing and then injected into embryos for phiC31 insertion.

### Muscle preparations and immunofluorescence in Drosophila

Muscle preparations in pupal abdomens were performed as previously described (Ribeiro et al., 2011). Staged pupae were removed from the pupal case and pinned on a sylgard-covered petri dish in dissection buffer (5 mM HEPES, 128 mM NaCl, 2 mM KCl, 4 mM MgCl_2_, 36 mM sucrose, pH 7.2). Abdomens were opened with microscissors, pinned flat, and fixed at room temp for 20 min. (4% formaldehyde, 50 mM EGTA, PBS). Then, the samples were unpinned and blocked at room temp for 30 min (0.3% bovine serum albumin, 2% goat serum, 0.6% Triton X100, PBS), incubated with primary antibody overnight at 4°C, washed (0.1% Triton PBS), then incubated for 2 h at room temp with Alexa Fluor488-conjugated secondary antibodies (Thermo Fisher Scientific, Waltham, MA) and counterstained with phalloidin for F-actin as needed. The stained samples were washed and mounted in FluorSave reagent (Merck Millipore, Burlington, MA).

### Confocal fluorescence microscopy

For imaging of live pupal DIOMs, staged pupae were removed from the pupal case, mounted between slide-glass and cover-glass following a protocol (Zitserman and Roegiers, 2011), and imaged through the cuticle from the dorsal side. Live DIOMs were observed on a confocal microscope FV1000D with a 60x oil/1.35 NA UPlanSApo (Olympus, Tokyo, Japan) or FV3000 with a 60x silicone/1.30 NA UPlanSApo (Olympus, Tokyo, Japan). The image acquisition software used was Fluoview (Olympus, Tokyo, Japan). The exported images were adjusted and analyzed using the ImageJ.

### Electron microscopy

Staged pupae (20 h or 4 d APF) were removed from pupal cases, pinned on a sylgard-covered petri dish, dissected directly in fixative (2% paraformaldehyde, 2.5% glutaraldehyde, 150 mM sodium cacodylate, pH 7.4) and fixed for 2 h at room temp and then overnight at 4°C. The dissected fillets were washed with 0.1 M phosphate buffer pH 7.4, post-fixed in 1% OsO_4_ buffered with 0.1 M phosphate buffer for 2 h, dehydrated in a graded series of ethanol, and embedded flat in Epon 812. Ultrathin sections, the thickness of 70 nm, were collected on copper grids covered with Formvar, double-stained with uranyl acetate and lead citrate, and then observed by a transmission electron microscope, H-7100 (Hitachi, Tokyo, Japan).

### Spinning-disk time-lapse imaging

Twenty h APF DIOMs expressing GFP:Atg8 and Spin:RFP were imaged every 60 s for 30 min using a 60x silicone/1.30 NA UPlanSApo objective (Olympus, Tokyo, Japan) on an inverted microscope (IX83; Olympus, Tokyo, Japan) with a spinning-disc confocal scanner unit (Dragonfly; Andor Technology, Belfast, UK) and a CMOS camera (Zyla 4.2; Andor Technology, Belfast, UK). Z-series images of each time point were exported using Fusion (Andor Technology, Belfast, UK). Then, the exported files were cropped, thresholded, and analyzed by the ImageJ.

### FRAP analysis

Twenty-four h APF DIOMs expressing both Cp1:mKO and GFP:Stx17 were imaged using a 60x silicone/1.30 NA UPlanSApo objective on a confocal microscope, FV3000 (Olympus, Tokyo, Japan). Three frames were acquired before photobleaching. Bleaching was performed on equally sized rectangular regions of interest (ROI) with a 568-nm laser at 30.7% power for 2 ms per pixel. Images of each channel were immediately acquired after the bleaching at a 20 s interval for 440 s. ROI mean intensities were measured using the ImageJ. FRAP analysis to determine normalized fluorescence intensity was performed as described (Goodwin and Kenworthy, 2005).

### Fly injection

Micro-injection was carried out using a stereomicroscope (SZX16; Olympus, Tokyo, Japan) with a manipulator (M-152; Narishige, Tokyo, Japan), an injector (IM-400; Narishige, Tokyo, Japan), and a compressor (0.2LE-8SB; Hitachi, Tokyo, Japan). LysoTracker Red was diluted in DMSO to 1 mM, and DQ-BSA was diluted in water to 10 mg/mL. The micro-injection needle was loaded with the solutions and then introduced into the abdomen of staged pupae. Although we could not control the exact amount of the solution for each injection, because of limitations imposed by our injection system, we roughly estimate the amount is 5 to 10 nL. Injected pupae were cultured at 25°C for 15 min for LysoTracker or 3 h for DQ-BSA, and observed on FV3000.

### Image analyses

Quantification of tubules was performed as follows: the percentage of DIOMs containing tubule >5 µm in length was manually counted. At least 8 DIOMs were checked in each animal. More than 10 animals were analyzed in each genotype (Fig. 2I, 3A-J, S2D, and S3A-E). Alternatively, the total tubule area (i.e., the sum of the area of all tubules) per unit area was measured by using the ImageJ. Intensities were binarized and then skeletonized by LpxLineExtract, which is invoked by Lpx_Filter2d plugin in the LPixel ImageJ plugins package (Kuki et al., 2017). The >5 µm skeletonized lines in the images were extracted by using the analyze particles function in Fiji. More than 10 cropped images were used for the quantification. The total tubule area per unit area was shown as an arbitrary value (Fig. 1E and S1D).

The distance between points at which GFP:Atg8 puncta disappeared or randomly simulated puncta and the nearest Spin:RFP-positive tubule was measured by using the measure function in the ImageJ (Fig. 2G). The Spin:RFP-positive tubules in the images were extracted, as mentioned above. As a simulation, we randomly drew puncta in images, in which GFP:Atg8-positive puncta disappeared, using the ImageJ plugin. More than 30 cases were used for the quantification.

Quantification of the diameter of vacuoles in Fig. 3L was performed as follows: the mCherry:GFP:Stx17-positive vacuole size was determined by a morphometric analysis using the ImageJ. More than 100 vacuoles from at least 10 DIOM images were analyzed for each genotype. Images of GFP-channel were used for the quantification.

To categorize DIOM remodeling phenotypes, 4 d APF animals were dissected and stained for Dlg1 and F-actin. More than 10 animals were analyzed for each genotype. At least 10 DIOMs per animal were observed and categorized into these four groups. 1) Regular, straight DIOM with organized myofibrils and T-tubules; 2) Thin myofibril layer, straight DIOM with thin myofibrils; 3) Disorganized, irregular shaped DIOM with disorganized myofibrils and T-tubules; 4) Detached, detached and rounded DIOM (Fig. 5D).

Quantification of the aspect ratio of DIOMs was performed as follows; 4 d APF animals expressing mCD8:GFP in muscle were observed by a fluorescent stereo microscope, SZX16 (Olympus, Tokyo, Japan) with a CMOS camera, ORCA-ER (Hamamatsu Photonics, Hamamatsu City, Japan). More than 50 DIOMs from at least five animals were analyzed each genotype. The aspect ratio, length per width, was determined by a morphometric analysis using the ImageJ (Fig. 5B, S2B, and S2F).

For the comparative analysis LysoTracker Red and DQ-BSA intensities, the acquired raw images were binarized, extracted, and measured the average fluorescent intensity of objects using the Fiji. The median intensity was set as one in each image. More than 250 objects from at least five images were analyzed for each time point.

### Statistics

Each experiment was performed at least three times as biological and technical replicates (at least three different cohorts of unique flies were analyzed in repeat procedures performed on at least three different days). One exception was for TEM analyses, which were performed on two parallel replicates with multiple animals each. All replicate experiments were performed in parallel with wild-type controls. The SD was used as error bars for bar charts from the mean value of the data. When more than two genotypes or treatments were used in an experiment, the statistical analysis was performed using Tukey’s test or Dunnett’s test on Prism8 software. An unpaired 2-tailed student’s t-test was used to compare two means. *F*-test was used to compare the dispersion between the two groups. p<0.05 was regarded as statistically significant. p<0.05 is indicated with single asterisks, and p<0.001 is indicated with double asterisks.

## Acknowledgments

We are grateful to GC. Chen (Academia Sinica), E. Kuranaga (Tohoku Univ.), JH. Lee (Univ. of Michigan), G. Davis (UCSF), T. Neufeld (Univ. of Minnesota), H. Stenmark (Oslo Univ. Hospital), Bloomington Drosophila Stock Center, DGRC, VDRC, FlyTrap, Kyoto DGGR, and NIG fly for reagents. We are grateful to members of the Fujita lab and Fukuda lab for helpful comments. This work was supported in part by Grant-in-Aid for Scientific Research (C) from the MEXT (grant number 18K06202 to NF), Grant-in-Aid for Scientific Research (B) from the MEXT (grant number 19H03220 to MF), Japan Science and Technology Agency (JST) PRESTO (grant number JPMJPR18H8 to NF), JST CREST (grant Number JPMJCR17H4 to MF), and the research grant of Astellas Foundation for Research on Metabolic Disorders (to NF).

## Abbreviations used

ALR: autophagic lysosome reformation
APF: after puparium formation
ATG: autophagy-related
Cp1: cysteine protease 1
DIOM: dorsal internal oblique muscle
DQ: dye quenched
FRAP: fluorescence recovery after photobleaching
mTOR: mechanistic target of rapamycin
SNARE: soluble NSF attachment protein receptor
Stx17: Syntaxin17
tAL: tubular autolysosome
TEM: transmission electron microscopy
V-ATPase: vacuolar H^+^ ATPase
3IL: third instar larvae.

## Author contributions

NF designed the research; TM, YK, and NF performed the experiments; TM, YS, AAK, MF, and NF analyzed and interpreted data; NF took the lead in writing the manuscript. All authors provided critical feedback and helped shape the research and manuscript.

## Competing financial interests

The authors declare no competing financial interests.

## Supplementary figure legends

**Figure 2–figure Supplement 1.**
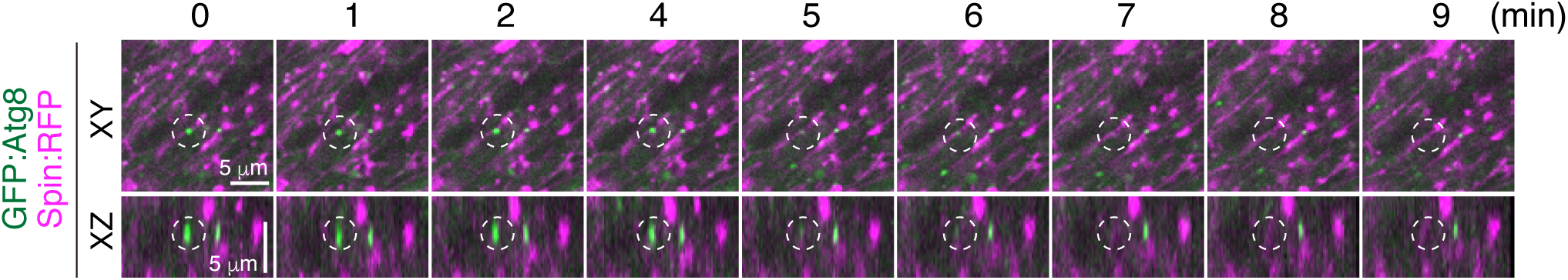
Time-lapse imaging of Spin:RFP and GFP:Atg8a in 20 h APF DIOMs. Top row, XY slices. Bottom row, XZ slices.

**Figure 2–figure Supplement 2.**
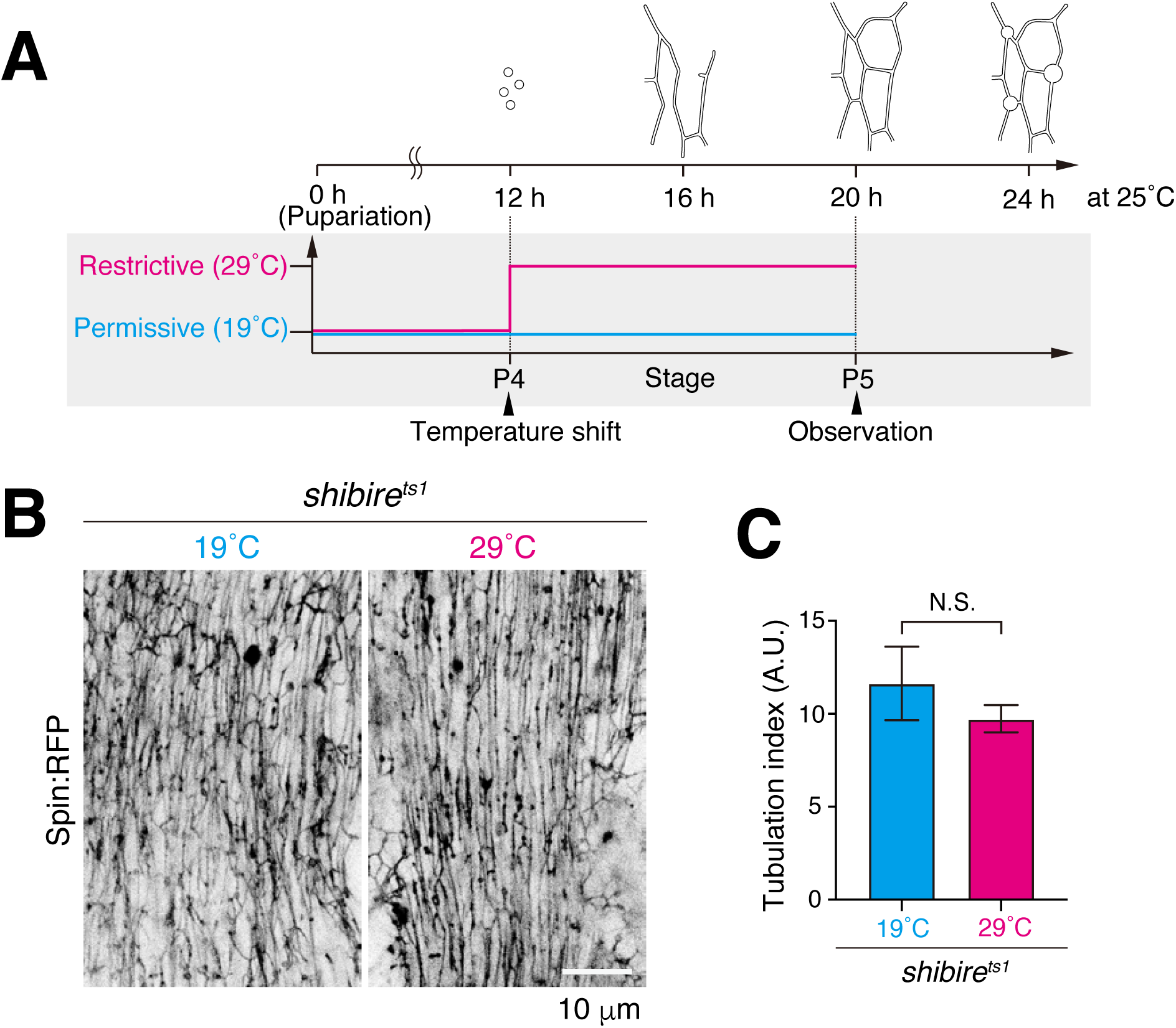
Blockade of *shibire* on tAL network formation (A) Scheme of an experiment using *shibire* temperature-sensitive mutant (*shi*^*ts1*^). *shi*^*ts1*^ mutant was incubated at 19°C all the time (Permissive) or 29°C from P4 to P5 stage (Restrictive). (B and C) Spin:RFP-positive tubular network in DIOM at P5 stage (B). Quantification of the Spin:RFP-positive tubules in panel B (C). The average ± SD is shown, n=5. NS, not significant; (Student’s *t*-test).

**Figure 2–figure Supplement 3.**
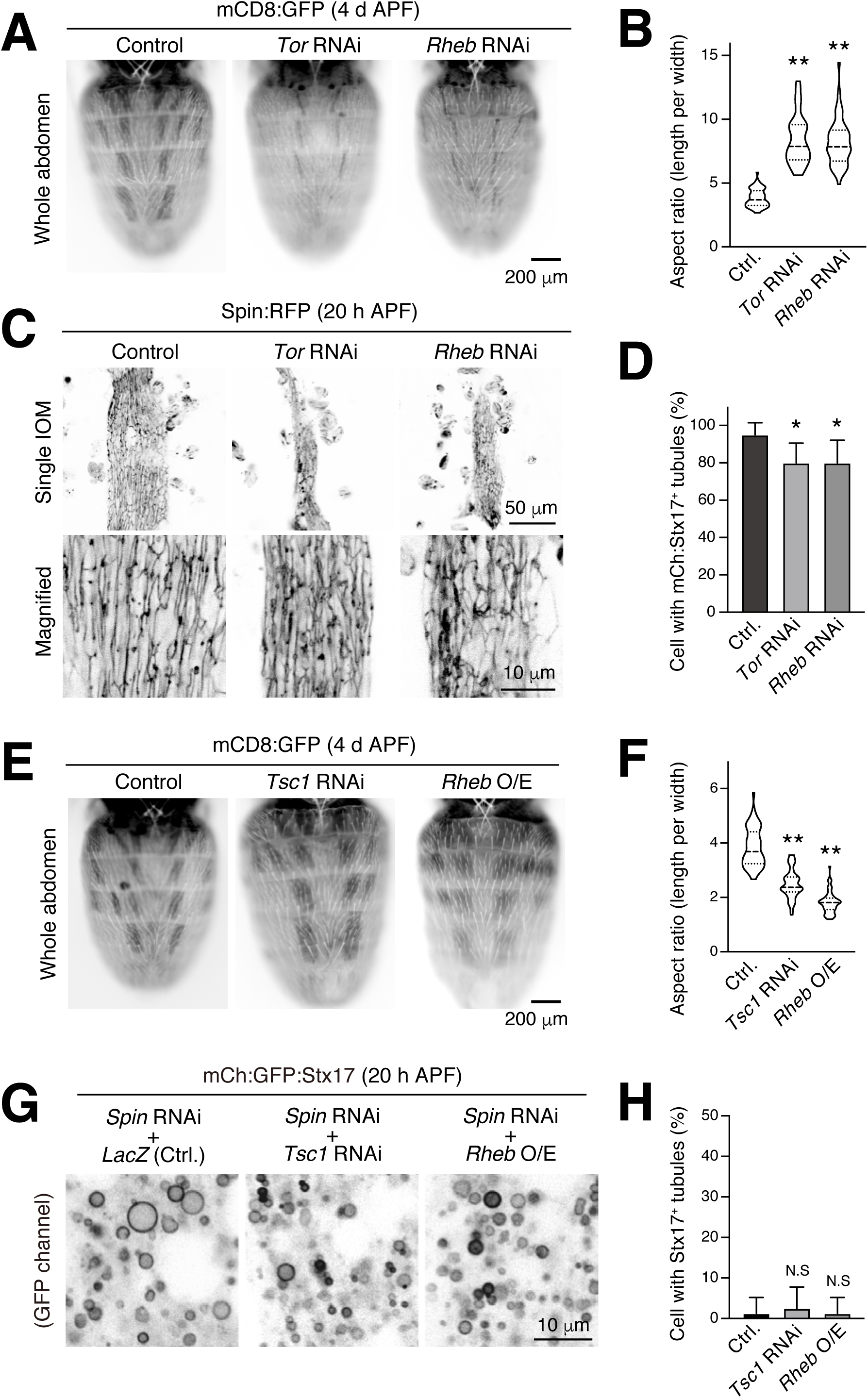
Forced activation or inactivation of mTOR in DIOMs (A and B) Effect of forced mTOR inactivation on the shape of DIOMs at 4 d APF (A). Violin plot of the aspect ratio of DIOMs, n>40. **, p<0.001 (Dunnett’s test) (B). (C and D) Effect of forced mTOR inactivation on the formation of the Spin:RFP-positive tubular structures in 20 h APF DIOMs (C). Mean percent + SD of DIOMs with Spin:RFP-positive tubules, n=10. *, p<0.05 (Dunnett’s test) (D). (E and F) Effect of forced mTOR activation on the shape of DIOMs at 4 d APF (E). Violin plot of the aspect ratio of DIOMs, n>50. **, p<0.001 (Dunnett’s test) (F). (G and H) Genetic interaction between *Spin* and mTOR regulators. Combination of *Spin* RNAi and *Tsc1* RNAi or *Rheb* overexpression on mCherry:GFP:Stx17-positive structures in 20 h APF DIOMs (G). Mean percent + SD of DIOMs with Stx17-positive tubules, n=10. N.S, not significant (Dunnett’s test) (H).

**Figure 3–figure Supplement 1.**
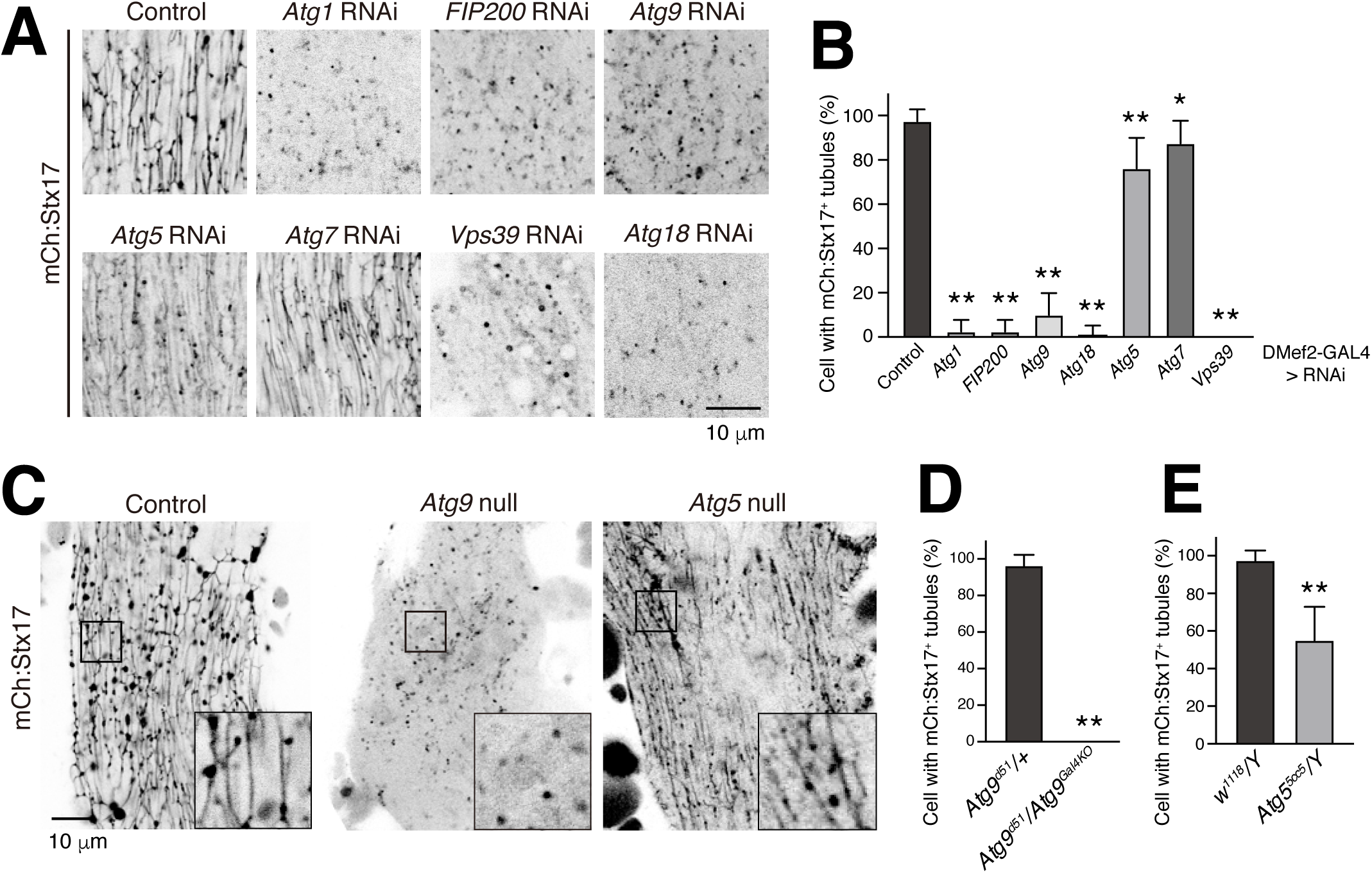
Loss of *ATG* genes on mCh:Stx17-positive tubules (A and B) Effect of RNAi of autophagy-related genes on mCherry:Stx17-positive tubules in 20 h APF DIOMs (A). Mean percent + SD of DIOMs with more than 5 μm mCherry:Stx17-positive tubules, n=10. *, p<0.05; **, p<0.001 (Dunnett’s test) (B). (C-E) mCherry:Stx17 in control, *Atg9* null, or *Atg5* null DIOMs at 20 h APF (C). Mean percent + SD of DIOMs with more than 5 μm mCherry:Stx17-positive tubules in control or *Atg9* null, n=10 (D), control or *Atg5* null, n=10 (E). **p<0.001 (Student’s *t*-test).

**Figure 3–figure Supplement 2.**
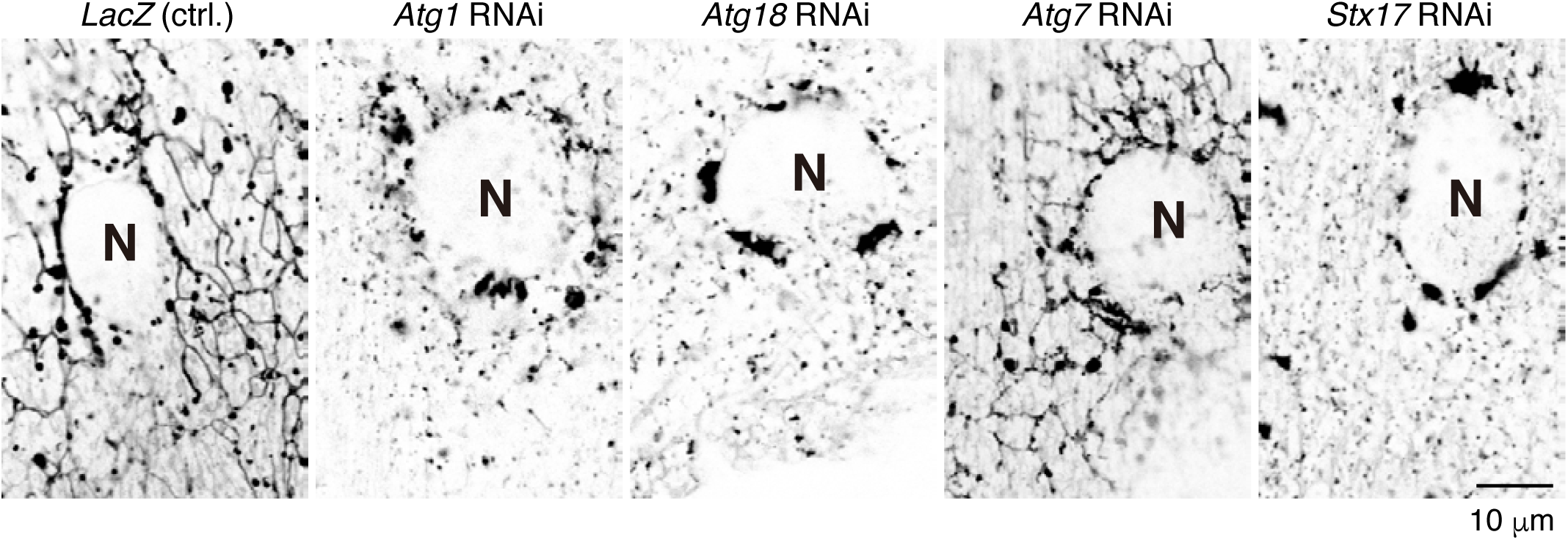
Effect of RNAi of autophagy-related genes on Spin:RFP-positive tubules in 3IL body wall muscle. The images are sections close to the sarcolemma. N, nucleus.

**Figure 4–figure Supplement 1.**
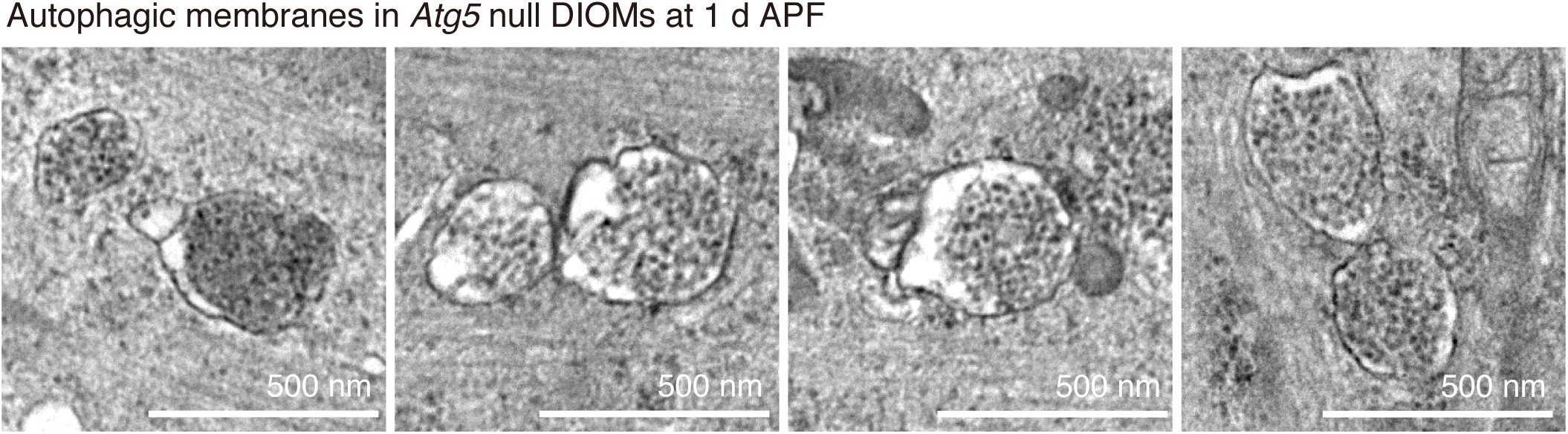
Typical double-membraned structures in the electron micrograms of *Atg5* null DIOMs at 20 h APF.

**Figure 5–figure Supplement 1.**
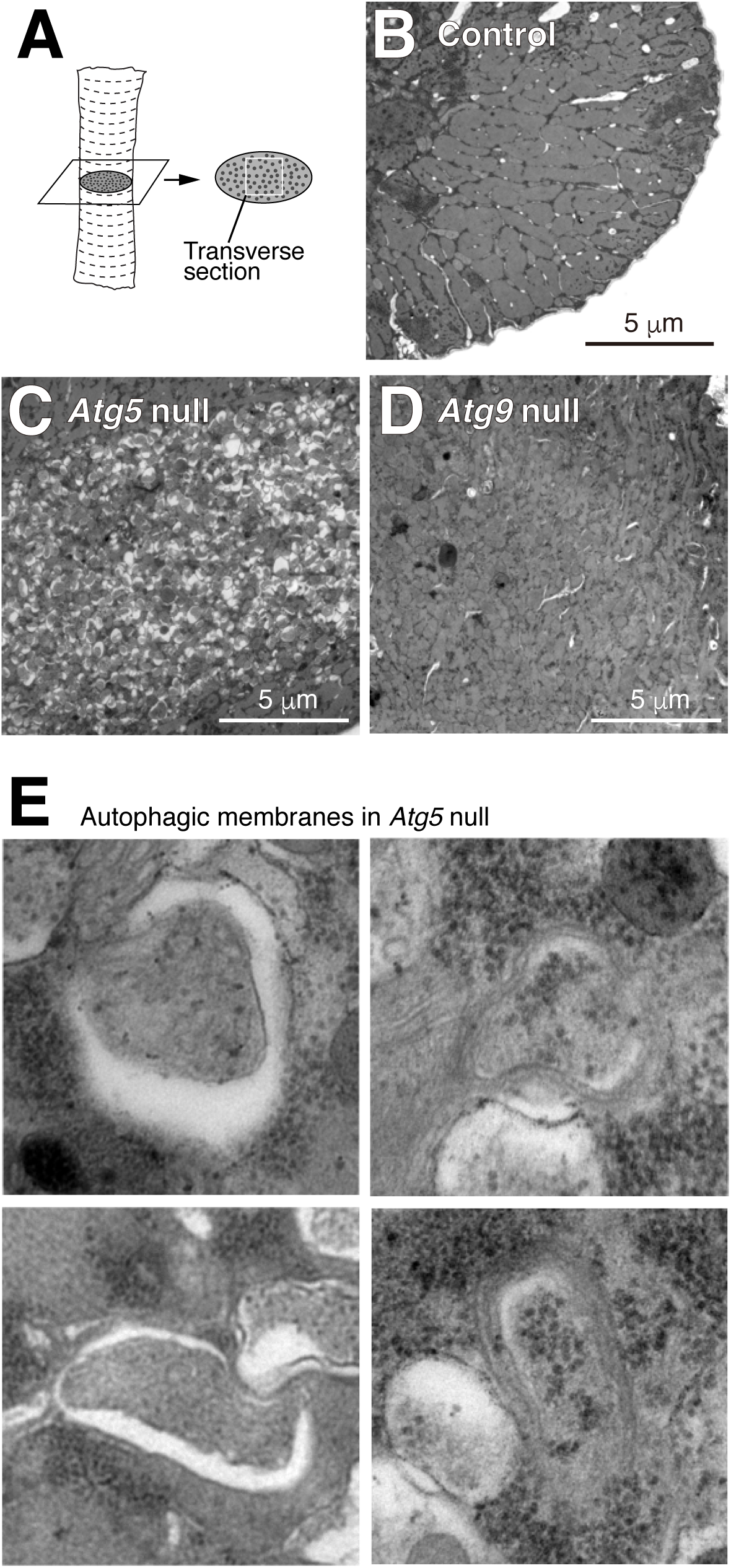
TEM analysis of *Atg5* null and *Atg9* null DIOMs at 4 d APF (A) Schematic of an DIOM TEM transverse section. (B-D) TEM images of 4 d APF DIOM transverse-sections of control (B), *Atg5* null (C), or *Atg9* null (D). (E) Typical examples of autophagic structures in *Atg5* null DIOMs at 4 d APF.

**Figure 7–figure Supplement 1.**
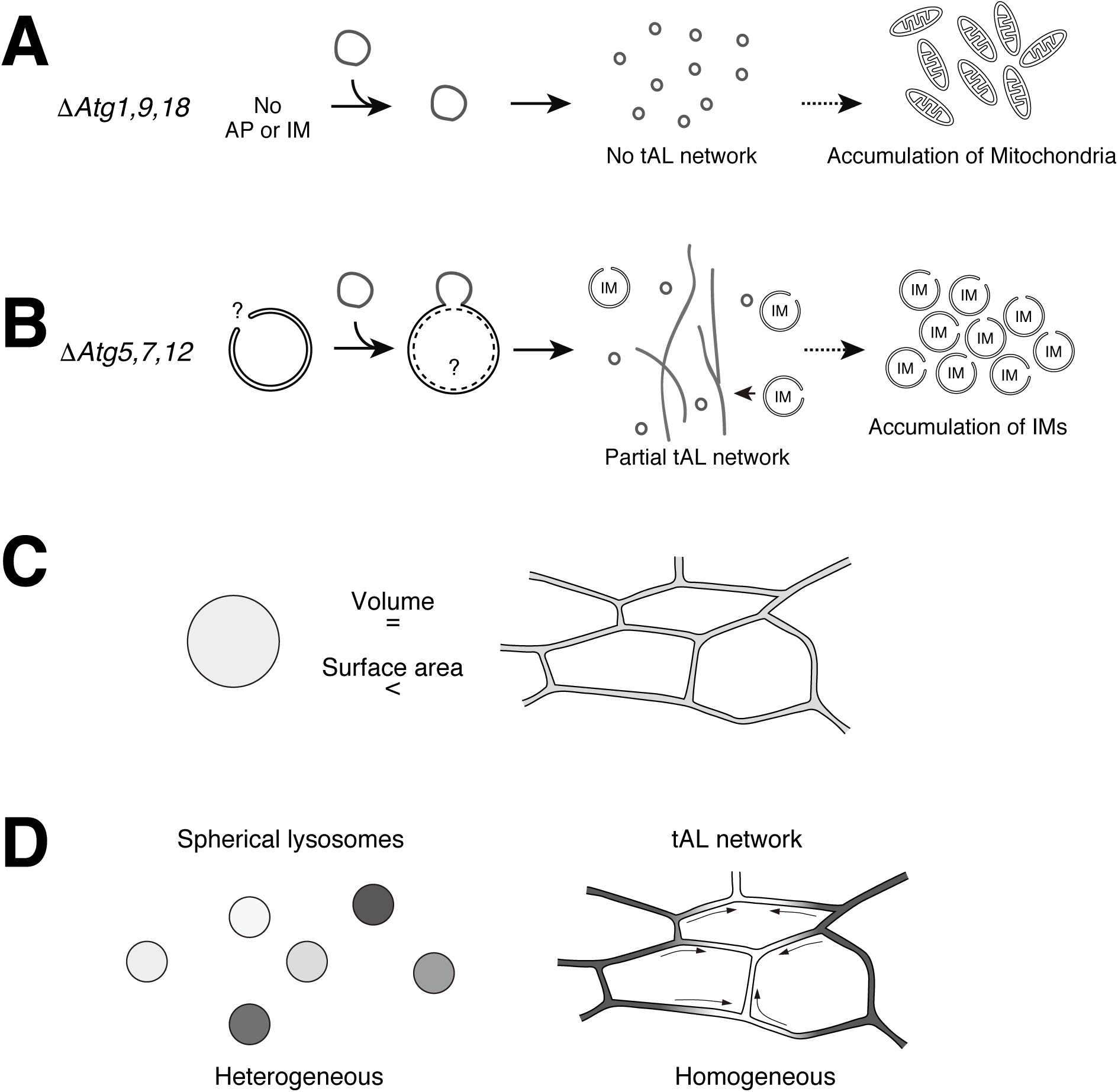
Possible model of the tAL network formation and its significance. (A) No autophagic membrane in loss of *Atg1, 9*, or *18*, which results in no tAL network. (B) Fusion of late isolation membranes with lysosomes leads to partially defective tAL network in loss of *Atg5, 7*, or *12*. (C) A ratio of surface area per volume of a ranging between 50 to 70-nm-diameter tube and 500-nm-diameter spherical vesicle. The tube has ∼5 times higher score than that of the vesicle. (D) The tAL network is homogeneous over a wide range compared to a number of spherical, discontinuous lysosomes.

## Supplementary file 1

Detailed *Drosophila* genotypes shown in figures.

**Figure 2–Supplementary video 1**

Time-lapse imaging of Spin:RFP and GFP:Atg8a in 20 h APF DIOMs. The DIOMs was imaged every 60 s for 30 min on an inverted microscope with a spinning-disc confocal scanner unit and a CMOS camera. Z-stacked images were shown.

## Source data file legends

**Figure 1-source data 1.** Relates to Figure 1E. Quantification of GFP:Stx17-positive tubules in DIOMs from 12 to 24 h APF (.xlsx file).

**Figure 2-source data 1.** Relates to Figure 2G. Quantification of the distance of Spin:RFP-positive tubule, and the point at which GFP:Atg8 puncta disappeared (GFP:Atg8) or randomly drawn puncta (random) (.xlsx file).

**Figure 2-source data 2.** Relates to Figure 2I. Quantification of the percentage of 20 h APF DIOMs with more than 5 μm GFP:Stx17-positive tubules in control, *Spin* RNAi, *TRPML* RNAi, or *Vha68-3* RNAi (.xlsx file).

**Figure 2-source data 3.** Relates to Figure 2–figure supplement 2C. Quantification of Spin:RFP-positive tubules in *shibire* temperature-sensitive mutant at 19°C all the time or 29°C from P4 to P5 stage (.xlsx file).

**Figure 2-source data 4.** Relates to Figure 2–figure supplement 3B. Quantification of the aspect ratio of DIOMs at 4 d APF of control, *Tor* RNAi, or *Rheb* RNAi (.xlsx file).

**Figure 2-source data 5.** Relates to Figure 2–figure supplement 3D. Quantification of the percentage of 20 h APF DIOMs with more than 5 μm Spin:RFP-positive tubules in control, *Tor* RNAi, or *Rheb* RNAi (.xlsx file).

**Figure 2-source data 6.** Relates to Figure 2–figure supplement 3F. Quantification of the aspect ratio of DIOMs at 4 d APF of control, *Tsc1* RNAi, or *Rheb* O/E (.xlsx file).

**Figure 2-source data 7.** Relates to Figure 2–figure supplement 3H. Quantification of the percentage of 20 h APF DIOMs with more than 5 μm Spin:RFP-positive tubules in control, *Tsc1* RNAi, or *Rheb* O/E (.xlsx file).

**Figure 3-source data 1.** Relates to Figure 3B. Quantification of the percentage of 20 h APF DIOMs with more than 5 μm Spin:RFP-positive tubules in *ATG* RNAi conditions (.xlsx file).

**Figure 3-source data 2.** Relates to Figure 3F. Quantification of the percentage of 20 h APF DIOMs with more than 5 μm Spin:RFP-positive tubules in control or *Atg9* null (.xlsx file).

**Figure 3-source data 3.** Relates to Figure 3H. Quantification of the percentage of 20 h APF DIOMs with more than 5 μm Spin:RFP-positive tubules in control or *Stx17* null (.xlsx file).

**Figure 3-source data 4.** Relates to Figure 3J. Quantification of the percentage of 20 h APF DIOMs with more than 5 μm Spin:RFP-positive tubules in control, *Atg5* null, or combination of *Atg5* null and *Stx17* RNAi (.xlsx file).

**Figure 3-source data 5.** Relates to Figure 3L. Quantification of the diameter of mCherry:GFP:Stx17-positive vesicles in co-RNAi of *Spin* and *Atg18* or *Vps39* (.xlsx file).

**Figure 3-source data 6.** Relates to Figure 3–figure supplement 1B. Quantification of the percentage of 20 h APF DIOMs with more than 5 μm mCh:Stx17-positive tubules in *ATG* RNAi conditions (.xlsx file).

**Figure 3-source data 7.** Relates to Figure 3–figure supplement 1D. Quantification of the percentage of DIOMs with more than 5 μm mCh:Stx17-positive tubules in control or *Atg9* null (.xlsx file).

**Figure 3-source data 8.** Relates to Figure 3–figure supplement 1E. Quantification of the percentage of 20 h APF DIOMs with more than 5 μm mCh:Stx17-positive tubules in control or *Atg5* null (.xlsx file).

**Figure 5-source data 1.** Relates to Figure 5B. Quantification of the aspect ratio of DIOMs at 4 d APF in *ATG* RNAi conditions (.xlsx file).

**Figure 5-source data 2.** Relates to Figure 5D. Quantification of the DIOM phenotypes at 4d APF in control, *Atg5* null, or *Atg9* null DIOMs (.xlsx file).

**Figure 6-source data 1.** Relates to Figure 6E. Quantification of the recovery of Cp1:mKO intensity after the bleaching (.xlsx file).

**Figure 7-source data 1.** Relates to Figure 7B. Quantification of the intensity of LysoTracker-positive objects in 12 or 24 h APF DIOMs (.xlsx file).

**Figure 7-source data 2.** Relates to Figure 7D. Quantification of the intensity of DQ Red-BSA-positive objects in 12 or 24 h APF DIOMs (.xlsx file).

